# Comparison of multiple tractography methods for reconstruction of the retinogeniculate visual pathway using diffusion MRI

**DOI:** 10.1101/2020.09.19.304758

**Authors:** Jianzhong He, Fan Zhang, Guoqiang Xie, Shun Yao, Yuanjing Feng, Dhiego C. A. Bastos, Yogesh Rathi, Nikos Makris, Ron Kikinis, Alexandra J. Golby, Lauren J. O’Donnell

## Abstract

The retinogeniculate visual pathway (RGVP) conveys visual information from the retina to the lateral geniculate nucleus. The RGVP has four subdivisions, including two decussating and two non-decussating pathways that cannot be identified on conventional structural magnetic resonance imaging (MRI). Diffusion MRI tractography has the potential to trace these subdivisions and is increasingly used to study the RGVP. However, it is not yet known which fiber tracking strategy is most suitable for RGVP reconstruction. In this study, four tractography methods are compared, including constrained spherical deconvolution (CSD) based probabilistic (iFOD1) and deterministic (SD-Stream) methods, and multi-fiber (UKF-2T) and single-fiber (UKF-1T) unscented Kalman filter (UKF) methods. Experiments use diffusion MRI data from 57 subjects in the Human Connectome Project. The RGVP is identified using regions of interest created by two clinical experts. Quantitative anatomical measurements and expert anatomical judgment are used to assess the advantages and limitations of the four tractography methods. Overall, we conclude that UKF-2T and iFOD1 produce the best RGVP reconstruction results. The iFOD1 method can better quantitatively estimate the percentage of decussating fibers, while the UKF-2T method produces reconstructed RGVPs that are judged to better correspond to the known anatomy and have the highest spatial overlap across subjects. Overall, we find that it is challenging for current tractography methods to both accurately track RGVP fibers that correspond to known anatomy and produce an approximately correct percentage of decussating fibers. We suggest that future algorithm development for RGVP tractography should take consideration of both of these two points.

## 1. Introduction

The retinogeniculate pathway (RGVP) begins at the retina, passes through the optic nerve, the optic chiasm, and the optic tract, and finally connects to the lateral geniculate nucleus (LGN) (J. Salazar et al., 2019; Purves et al., 2001). See Figure 1 for an anatomical overview. Approximately 53-58 percent of RGVP fibers decussate (cross) at the optic chiasm (Chacko, 1948; Kupfer et al., 1967; v. Sántha, 1932). Therefore, the RGVP can be divided into four anatomical subdivisions: two decussating (medial) and two non-decussating (lateral) fiber pathways, and each subdivision undertakes the important task of conveying visual information of visual hemi-fields (Purves et al., 2014; Quigley et al., 1982; Quigley & Green, 1979).

**Figure 1.**
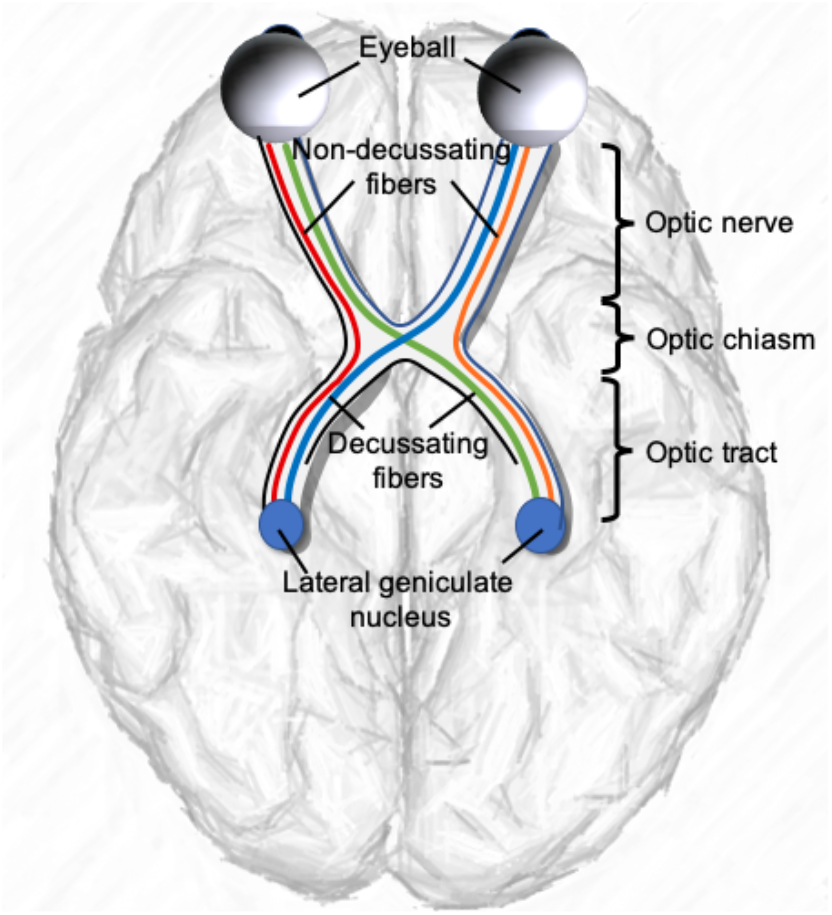
A schematic anatomical overview of the RGVP. There are four major subdivisions, including two non-decussating fiber pathways (orange and red) connecting the optic nerve and the optic tract in the same hemisphere and two decussating fiber pathways (green and blue) connecting the optic nerve and the optic tract across the hemispheres.

The study of the RGVP is important in multiple research and clinical applications. The RGVP can be affected in many diseases, including pituitary tumors (Laws et al., 1977; Rucker & Kernohan, 1954), glioma (Hales et al., 2018), optic neuritis (Beck et al., 2003; Brex et al., 2002; Optic Neuritis Study Group, 1997), ischemic optic neuropathy (Attyé et al., 2018; Cho et al., 2016), and optic nerve sheath meningioma (Schick et al., 2004; Turbin et al., 2002). In particular, visualization of the RGVP can be helpful in planning surgical approaches to lesions intrinsic or extrinsic to the pathway. Neoplasms of the sellar region, such as pituitary adenomas, tuberculum sellae meningioma, Rathke’s cleft cysts, and craniopharyngiomas can cause extrinsic compression of the RGVP fibers when extending superiorly (Ma et al., 2016). Optic pathway glioma (OPG), a childhood tumor most common in patients with neurofibromatosis type 1 (NF-1), is a diffusely infiltrative tumor with ill-defined boundaries, causing intrinsic compression and infiltration of the RGVP (Aquilina et al., 2015). Surgical resection of these tumors is challenging due to the high potential for damage to the visual apparatus. There is a poor correlation between tumor extent on conventional imaging modalities and visual impairment; therefore, detailed information about RGVP fiber pathway microstructure could potentially help clinicians in treatment decision making and surgical planning to potentially improve clinical outcomes (Aquilina et al., 2015; Fisher et al., 2012). The study of the complete RGVP is also relevant in the setting of high tension glaucoma, which can affect the whole optic pathway (Lestak et al., 2012).

Magnetic resonance imaging (MRI) techniques have been used to identify the RGVP for clinical and research purposes (Gala, 2015). Traditional T1-weighted (T1w) and T2-weighted (T2w) MRI are the most widely used, e. g, to compute the cross-sectional area of the intraorbital optic nerve (Gass et al., 1996; Simon J. Hickman, 2007; Simon J. Hickman et al., 2004; S. J. Hickman et al., 2001; Watanabe et al., 2008). However, these MRI sequences only indicate whether the RGVP is present at a particular location, while the 3D fiber pathway of the RGVP and the continuity of its subdivisions cannot be assessed. Diffusion MRI (dMRI), via a process called tractography, can track brain white matter and nerve fibers in vivo non-invasively based on the principle of detecting the random motion of water molecules in neural tissue (Basser et al., 1994, 2000). One advantage of dMRI is that it can enable tracking of the 3D trajectory of the RGVP for visualization of structures not visualized by traditional MRI sequences, e.g., crossing fibers at the optic chiasm (Puzniak et al., 2019). Multiple studies have investigated reconstruction of the RGVP using dMRI tractography (see Table 1) (Altintaş et al., 2017; Ather et al., 2019; Hofer et al., 2010; Panesar et al., 2019; Yoshino et al., 2016).

**Table 1.**
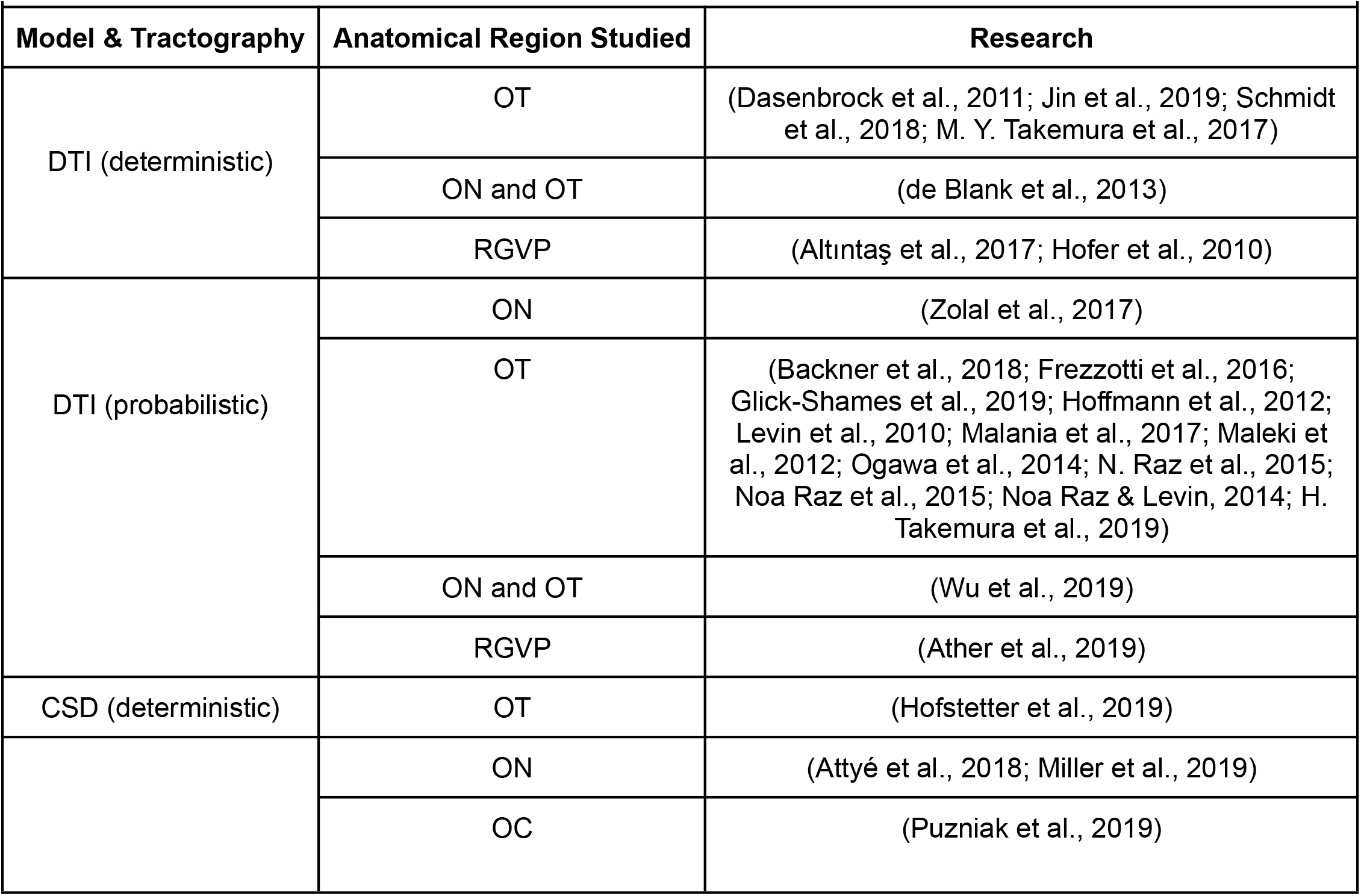
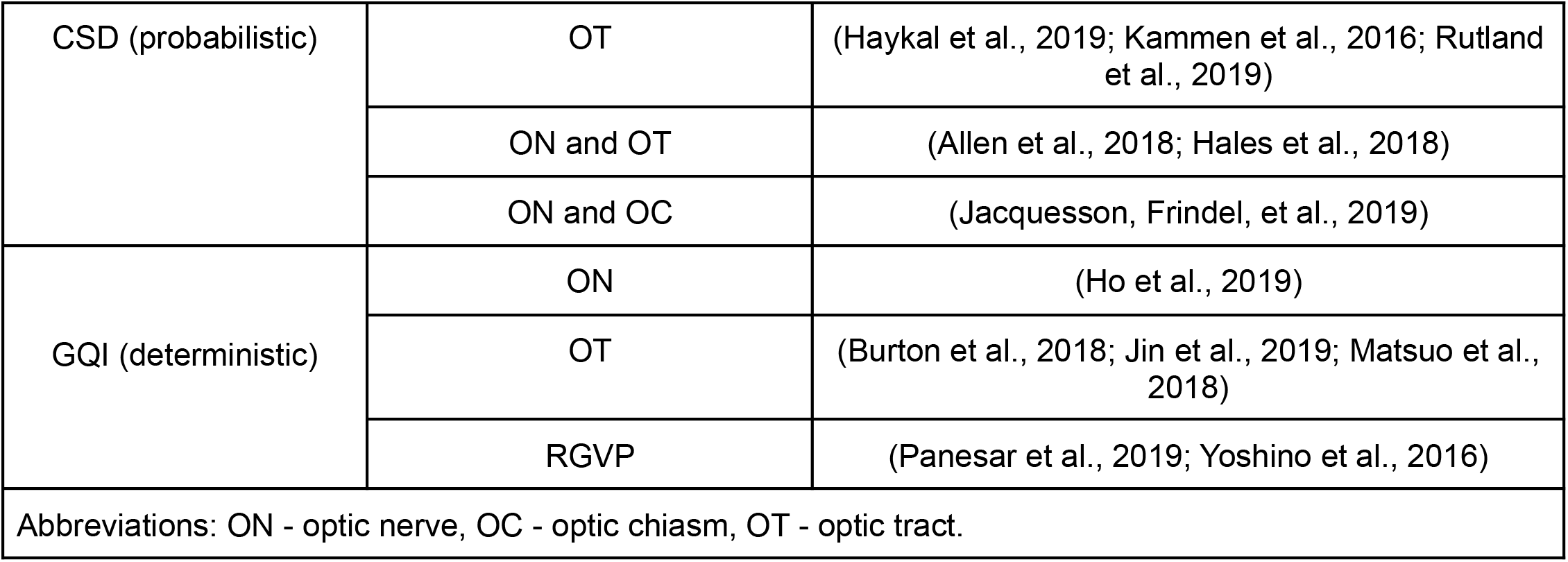
Summary of published tractography studies of the RGVP and its subregions, organized according to the fiber reconstruction methods employed and the anatomical region studied.

Despite the wide use of dMRI for RGVP reconstruction, practical questions remain about which tractography methods can provide a comprehensive delineation of the RGVP fiber pathway. The RGVP is located within the complex skull base environment, which contains nerve, bone, air, soft tissue, and cerebrospinal fluid. These anatomical structures can result in partial voluming, where voxels contain mixed information from multiple components, and susceptibility artifacts, which affect nerve fiber tracking performance. In addition, a unique characteristic of the RGVP is that there are crossing nerve fibers at the optic chiasm, which requires a tractography method to be highly sensitive despite the presence of crossing fibers. Recent reviews came to the overall conclusion that the choice of tractography methods and/or parameter settings strongly affects cranial nerve fiber reconstruction, including the RGVP (Jacquesson, Frindel, et al., 2019; Shapey et al., 2019). Table 1 gives a summary of existing tractography-based RGVP studies to describe the current state of the art in terms of the fiber tracking methods being used, among which the standard diffusion tensor imaging (DTI) tractography and more advanced methods such as those using constrained spherical deconvolution (CSD) and generalized q-sampling imaging (GQI) are most widely used. While these studies have shown highly promising RGVP tracking performance, most have focused on certain RGVP subregions of interest, e.g., the optic nerve, the optic tract, and/or the RGVP fibers within the optic chiasm (as summarized in Table 1), but not the complete RGVP pathway.

The main contribution of this work is to investigate, for the first time, the performance of multiple tractography methods for reconstruction of the complete RGVP including the four anatomical subdivisions. Four different tractography methods are compared, including two methods based on the CSD model and two methods that use the unscented Kalman filter (UKF) tractography framework. We choose the CSD-based tractography methods because the CSD model is currently the most widely applied method for cranial nerve fiber tracking (Table 1), and we choose the UKF methods because UKF has recently shown high performance in reconstructing the cranial nerves (Xie et al., 2020a; Zhang, Xie, et al., 2020). A comprehensive quantitative and visual comparison is performed using dMRI data from the Human Connectome Project (HCP) (Glasser et al., 2013; Van Essen et al., 2013). Quantitative evaluation includes the reconstruction rate of the four anatomical subdivisions, the percentage of decussating fibers, and a comparison to anatomical-T1w-based RGVP segmentation. Furthermore, to assess the reliability of the RGVP tracking results, an inter-expert validation is performed to compare the spatial overlap of RGVPs reconstructed from two experts. Next, following the popular strategy of assessing tractography using anatomical expert judgment (D. Q. Chen, Zhong, et al., 2016; Pujol et al., 2015; Xie et al., 2020b), we performed an expert judgment experiment to rank the anatomical appearance of the reconstructed RGVPs. Finally, we quantified the performance of each tractography method across subjects using the normalized overlap score (NOS) method (D. Q. Chen, Zhong, et al., 2016).

## 2. Materials and Methods

### 2.1 Evaluation datasets

We used a total of 100 dMRI datasets (the ‘‘100 Unrelated Subjects’’ release) (54 females and 46 males, age: 22 to 35 years old) from the HCP database (https://www.humanconnectome.org) (Van Essen et al., 2013) for experimental evaluation. The HCP data scanning protocol was approved by the local Institutional Review Board (IRB) at Washington University. The HCP database provides dMRI data that was acquired with a high-quality image acquisition protocol using a customized Connectome Siemens Skyra scanner and processed using a well-designed processing pipeline (Glasser et al., 2013) including motion correction, eddy current correction, EPI distortion correction, and co-registration between dMRI and T1w data. The dMRI acquisition parameters in HCP were: TR=5520 ms, TE=89.5 ms, FA=78°, voxel size=1.25×1.25×1.25 mm^3^, and FOV=210×180 mm^2^. A total of 288 images were acquired in each dMRI dataset, including 18 baseline images with a low diffusion weighting b = 5 s/mm^2^ and 270 diffusion weighted (DW) images evenly distributed at three shells of b = 1000/2000/3000 s/mm^2^. In our study, we used the single-shell b=1000 s/mm^2^ data, consisting of 90 DW images and 18 baseline images, to perform fiber tracking using each compared tractography method (see Section 2.2 for details). We chose single-shell b=1000 s/mm^2^ data because it is similar to clinical acquisition protocols as applied in many previous RGVP-related studies (Altintaş et al., 2017; Attyé et al., 2018; Backner et al., 2018; Burton et al., 2018; de Blank et al., 2013; Frezzotti et al., 2016; Glick-Shames et al., 2019; Hofer et al., 2010; Hofstetter et al., 2019; Jacquesson, Frindel, et al., 2019; Jin et al., 2019; Lober et al., 2012; Malania et al., 2017; Maleki et al., 2012; Ogawa et al., 2014; Noa Raz et al., 2015; Noa Raz & Levin, 2014; Schmidt et al., 2018; Wu et al., 2019). Furthermore, single-shell b=1000 s/mm^2^ data has been shown in our previous study to be more effective for identification of cranial nerves than higher b values (Xie et al., 2020a). In addition, we also used the anatomical T1w data (co-registered with the dMRI), on which the RGVP was more visually apparent than on dMRI, for facilitating creation of tractography seeding masks (Section 2.2.1) and RGVP tracking performance evaluation (Section 2.3.4). The T1w image acquisition parameters were: TR=2400 ms, TE=2.14 ms, and voxel size=0.7×0.7×0.7 mm^3^. More detailed information about the HCP data acquisition and preprocessing can be found in (Glasser et al., 2013).

To avoid potential effects on fiber tracking and experimental comparison, we performed a visual check of the dMRI data for each of the 100 subjects and we excluded any subjects whose dMRI data had incomplete RGVP coverage (as illustrated in Figure 2(a)) and/or abnormal signal that resulted in negative diffusion tensor eigenvalues (as illustrated in Figure 2(b)). For anonymization, face removal had been performed in the HCP imaging data using the algorithm as described in (Milchenko & Marcus, 2013). This process can cause a removal of dMRI data near the retina, such that only part of the RGVP is present in the data (see Figure 2(a) for an illustration in one example subject). We identified 38 subjects with incomplete RGVP coverage, which were excluded from our study. In addition, we excluded another 5 subjects with apparently abnormal signals in the skull base region where the RGVP passes through. In these subjects, we found many voxels with b=0 signals lower than the b=1000 signals, potentially due to Gibbs ringing effects and partial voluming issues (Veraart et al., 2016; Zhang, Ning, et al., 2019) resulting in negative diffusion tensor eigenvalues (see Figure 2(b) for an illustration in one example subject), which could prevent fiber tracking of the RGVP. Overall, data from 57 subjects (male/female: 16/41; Age: 22-36) was included in our study.

**Figure 2.**
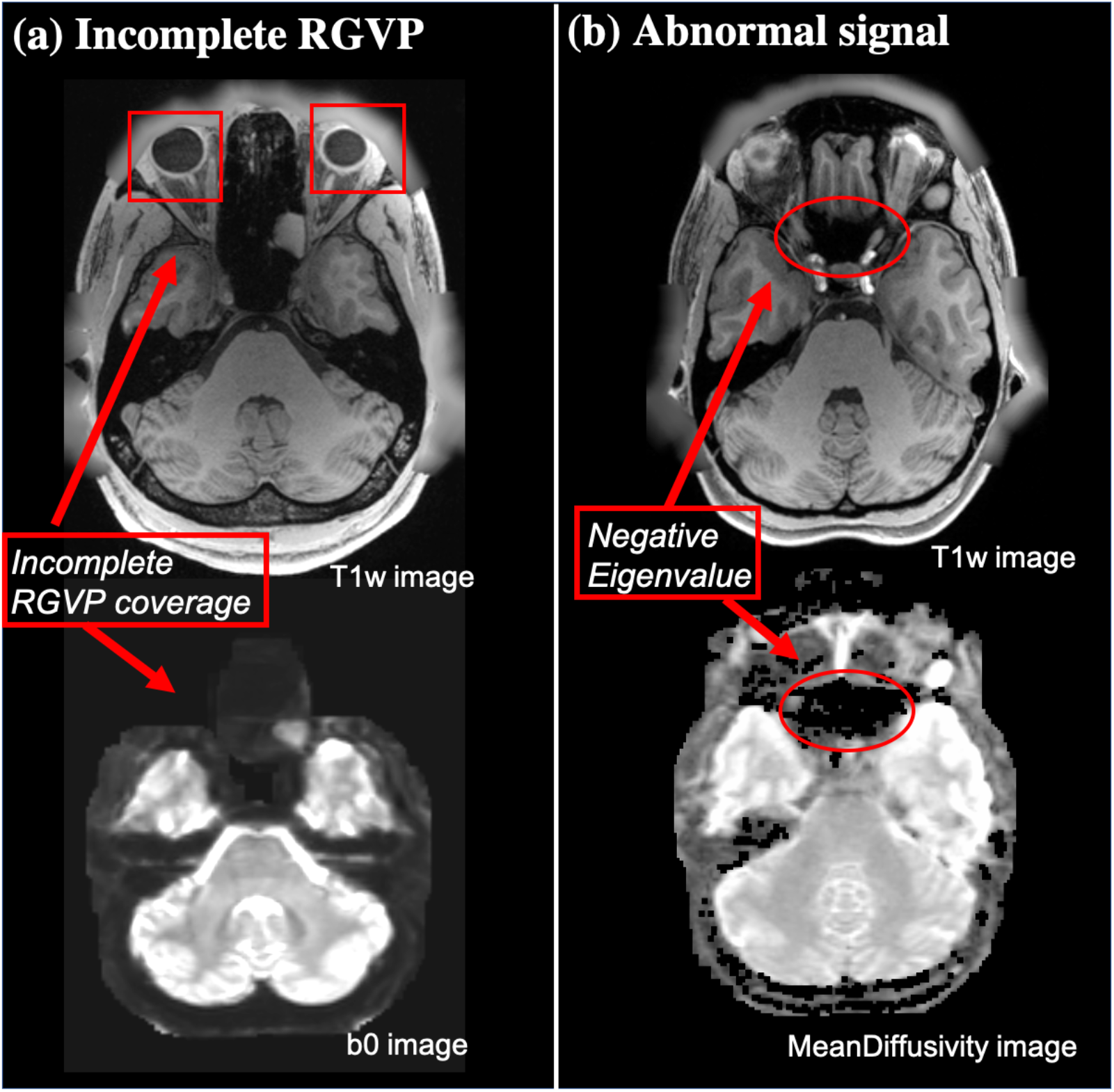
Illustration of two situations for excluded subjects. (a) shows an example of an incomplete RGVP in DWI data, where part of the optic nerve region is not present in the b0 image. (b) shows an example of abnormal signals, where the black holes on the mean diffusivity image show the voxels with negative diffusion tensor eigenvalues.

### 2.2 Comparison of four tractography methods

We compared four different tractography methods for reconstruction of the RGVP. The first two methods are *SD-Stream* and *iFOD1*, within the CSD tractography framework (J.-D. Tournier et al., 2004, 2007, 2012). We chose the CSD framework because it has been widely used in cranial nerve studies (Allen et al., 2018; Attyé et al., 2018; Hales et al., 2018; Haykal et al., 2019; Miller et al., 2019; Puzniak et al., 2019; Schmidt et al., 2018; Xie et al., 2020a). In brief, in the CSD-based tractography framework, fiber tracking works by stepping along a particular direction at each tracking step, with a constraint on the angle between successive steps. The direction for each step is obtained by sampling the fiber orientation distribution (FOD) at the current point. The SD-Stream method performs deterministic tracking where fiber tracking follows only the local direction of the FOD amplitude peak, whereas the iFOD performs probabilistic tracking by choosing a random direction of FOD, such that the probability of a particular direction being produced is proportional to the amplitude of the FOD along that direction. To achieve the best probabilistic tracking results, an initial experiment was performed to compare two CSD-based probabilistic methods, iFOD1 and iFOD2 (J. D. Tournier et al., 2010), over a large range of tracking parameter values (see Supplementary Material 1). Results indicated significantly better performance of iFOD1 on complete RGVP reconstruction, and therefore iFOD1 is used in the rest of the experiments in this paper. The other two compared methods are *UKF-1T* and *UKF-2T*, within the UKF tractography framework (Z. Chen et al., 2016; Gong et al., 2018; Liao et al., 2017; Malcolm et al., 2010; Reddy & Rathi, 2016; Zhang et al., 2018; Zhang, Xie, et al., 2020) has been shown to be highly effective in reconstructing cranial nerves (Xie et al., 2020a; Zhang, Xie, et al., 2020). It fits a tensor model to the diffusion data while tracking fibers, in a recursive estimation fashion (the current tracking estimate is guided by the previous one) using a Kalman filter. One benefit of the recursive estimation is to help stabilize model fitting; thus fiber tracking can be robust to a certain amount of imaging artifact/noise. In our study, we chose two types of UKF tractography. The UKF-1T method is a DTI tractography method, which fits a single tensor at each fiber tracking step, with the assumption that only one fiber passses through the current point. On the other hand, the UKF-2T method fits a mixture model of two tensors in which the first one represents the principal direction and the second one can represent crossing fibers.

#### 2.2.1 Tractography seeding

RGVP tractography was seeded (initiated) from all voxels within a mask, which was larger than the possible region through which the RGVP passes. This procedure enabled comprehensive RGVP reconstruction and was restricted to the potential RGVP region for efficiency, as applied in our previous cranial nerve tractography studies (Xie et al., 2020a; Zhang, Xie, et al., 2020). To boost the creation of the tractography seeding mask for all 57 subjects under study, we performed semi-automated mask processing, as illustrated in Figure 3(a). First, we manually drew a mask that covered the RGVP region on a T1w template (downloaded at: http://nist.mni.mcgill.ca/?p=858) in MNI space (Grabner et al., 2006). Then, we transferred this mask to each subject under study using a non-linear registration between the subject’s T1w data and the T1w template (FSL was used for computing the registration (Jenkinson et al., 2012)). In this way, the mask was transferred to each subject’s dMRI space that was co-registered with the T1w data. We visually checked the computed mask for each subject.

**Figure 3.**
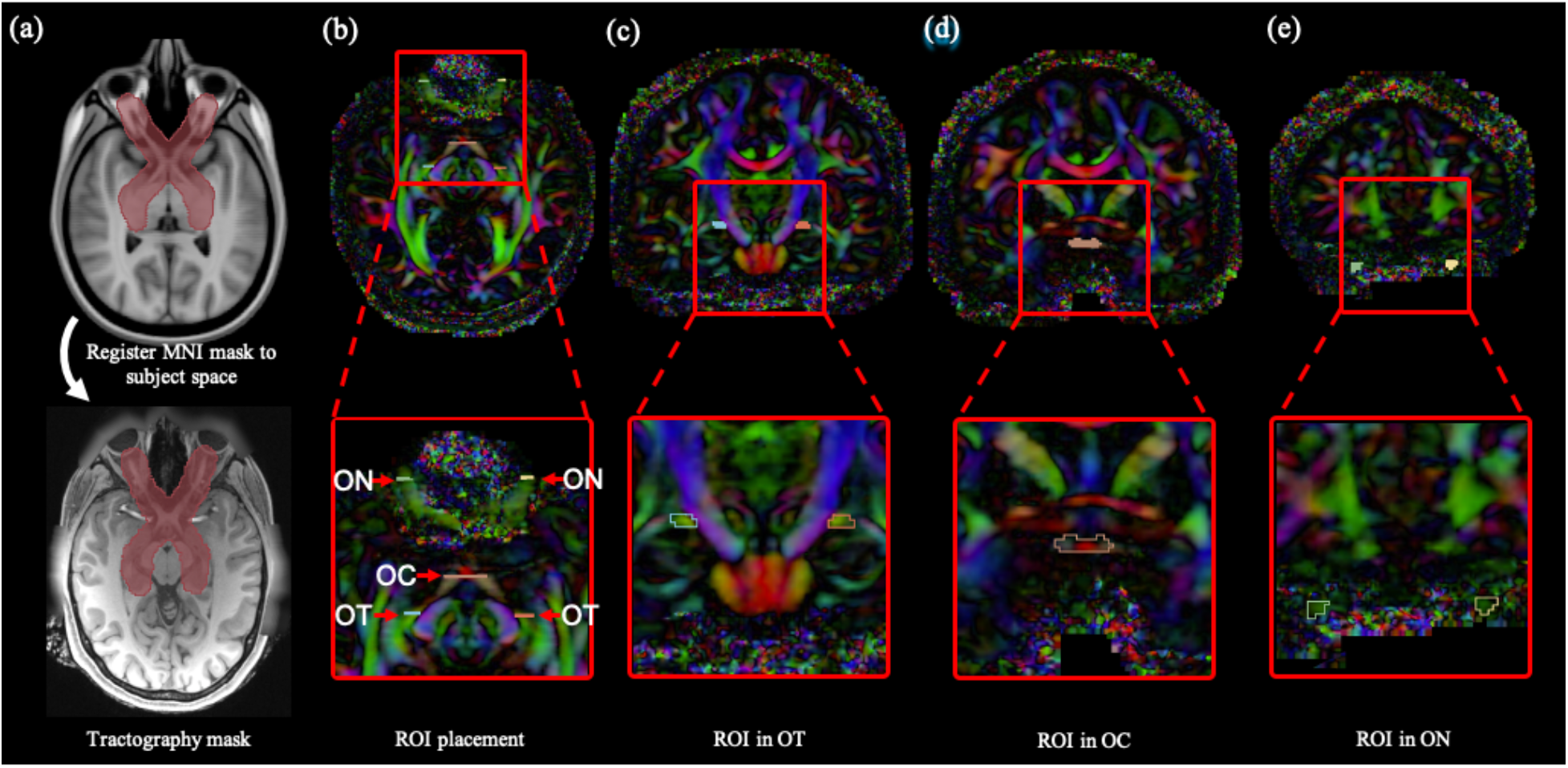
Tractography seeding mask and ROIs for selecting the RGVP fibers. (a) The tractography mask provides full coverage of the potential RGVP. (b) Five ROIs were drawn on the DTI image for every subject. (c) A pair of ROIs near the eyeball was drawn from the coronal view. (d) The second ROI was drawn at the optic chiasm. (e) The third ROIs were located in the optic tract near the LGN.

Following the creation of the seeding mask, RGVP fiber tracking was performed using each compared tractography method. To compare across the different methods, we first performed an experiment to determine the best-performing parameters for each tractography method. The goal of this experiment was to provide an unbiased comparison across methods by identifying method-specific parameters that provide good tracking results on single-shell b=1000 s/mm^2^ data, as in our previous work (Xie et al., 2020a). This step is critical because parameter settings will significantly influence the results of tractography (Parizel et al., 2007; H. Takemura et al., 2016; Wandell, 2016). Complete details for this experiment are provided in Supplementary Material 2. To briefly summarize, parameters affecting tractography starting and stopping were tested over a wide range of values. High-performing parameter combinations were identified according to their ability to track the four RGVP subdivisions. These final parameters for each tractography method are provided in Table 2. As expected, the identified seeding and stopping parameters are low to enable tracking through regions of low diffusion anisotropy at the skull base. For parameters related to stepwise tracking, the settings are similar to software defaults. For example, the maximum angle parameters of 10 degrees for iFOD1 and 80 degrees for SD-Stream are close to the defaults in MRtrix3 (15 degrees for iFOD1 and 60 degrees for SD-Stream). All other parameters of the tracking software packages were left as default. Overall, we generated the same number of fibers (n=40,000) using each tractography method, with a fiber length threshold of 45mm to eliminate any effect from fibers too short to form part of the RGVP anatomy. This length threshold is much lower than the actual length of the RGVP, where the axons of retinal ganglion cells are about 75mm long from eyeball to LGN (U. Schiefer & Hart, 2007). In our dataset, we verified this in several HCP subjects by measuring the distance from the eyeball to the LGN, following both ipsilateral and contralateral pathways.

**Table 2.** Best-performing parameters for each tractography method

| Table 2. Best-performing parameters for each tractography method |  |
| --- | --- |
| Tractography Methods | Tracking Parameters |
| SD-Stream | *seedCutoff*=0.006, *stopCutoff*=0.005, *maximumAngle*=80 |
| iFOD1 | *seedCutoff*=0.006, *stopCutoff*=0.005, *maximumAngle*=10 |
| UKF-1T | *SeedingFA*=0.02, *stoppingFA*=0.01, *Qm*=0.001, *Ql*=50 |
| UKF-2T | *SeedingFA*=0.02, *stoppingFA*=0.01, *Qm*=0.001, *Ql*=50 |

#### 2.2.2 ROI-based RGVP fiber selection

Next, we performed an ROI-based selection of the RGVP fibers from the seeded tractography data for each compared fiber tracking method. Five ROIs were used, including two from the anterior part of the bilateral optic nerves (near the eyeballs), one from the optic chiasm, and two from the posterior part of the bilateral optic tracts (near the LGN), as illustrated in Figure 3 (b-e). Our choice of the bilateral optic nerve ROIs and the optic chiasm ROI was because they have been widely used for selection of optic nerve fibers (Maleki et al., 2012; Miller et al., 2019; H. Takemura et al., 2019). For identifying the optic tract fibers, we chose to use ROIs in the optic tract near the LGN. Accurately delineating the LGN with the current MRI imaging resolution is a challenging task. Most of the existing studies that perform optic tract fiber selection use ROIs that cover (much larger than) the LGN, followed by further manual fiber filtering of false positive fibers (Allen et al., 2018; Haykal et al., 2019; Maleki et al., 2012). In our study, in order to avoid potential bias from further manual fiber selection, we chose ROIs in the optic tract that were easily identified in HCP imaging data. For each of the 57 subjects under study, the five ROIs were drawn by an clinical expert (G.X., who is a practicing neurosurgeon) on the directionally encoded color (DEC) map computed from the dMRI data (as shown in Figure 3 (b-e)). (Note that a second set of ROIs per subject were drawn by another clinical expert (S.Y., who is a neurosurgeon), which were used for inter-expert reliability validation, as in Section 2.3.4.) Drawing ROIs directly on the dMRI data has been shown to be effective for RGVP fiber selection (Ather et al., 2019).

Selection of the RGVP fibers was performed using the obtained ROIs. Specifically, for each RGVP subdivision, three ROIs were used, including one of the optic nerve ROIs (left or right), one of the optic tract ROIs (left or right), and the optic chiasm ROI. In total, four fiber selections were performed to obtain the four anatomical RGVP subdivisions, as shown in Figure 4. For the rest of the paper, we refer to these subdivisions as LL (non-decussating fibers connecting left optic nerve and left optic tract), RR (non-decussating fibers connecting right optic nerve and right optic tract), LR (decussating fibers connecting left optic nerve and right optic tract), and RL (decussating fibers connecting right optic nerve and left optic tract).

**Figure 4.**
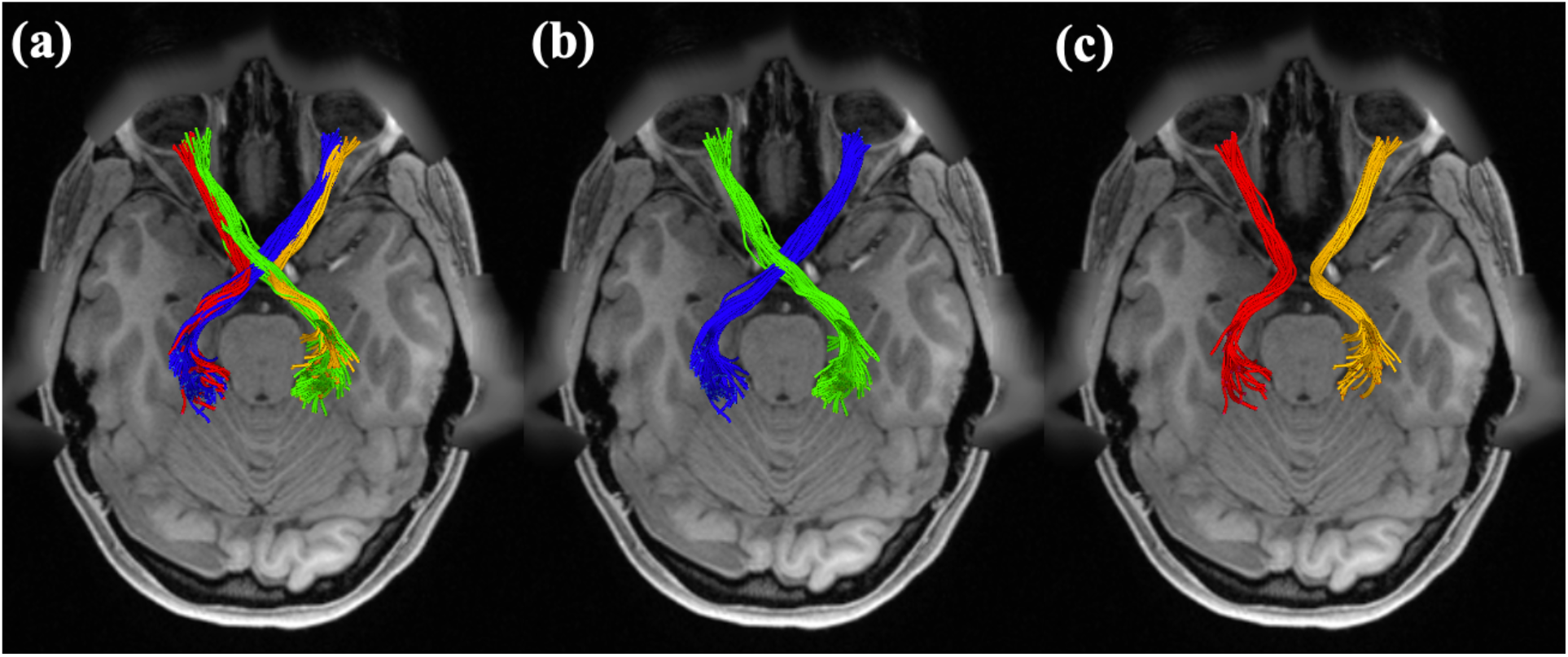
Illustration of the reconstructed subdivisions of the RGVP. (a) Axial view of an example RGVP overlaid on T1w image. Bundles of each color represent one subdivision. (b) A pair of decussating fiber bundles (green and blue), and (c) a pair of non-decussating fiber bundles (red and orange).

### 2.3 Experimental comparisons

We performed three experiments to assess the RGVP reconstruction performance of the different tractography methods. First, we compared the reconstruction rate of each RGVP subdivision across the methods (Section 2.3.1). Second, we computed the percentage of decussating fibers in the optic chiasm (Section 2.3.2), Third, we measured the correlation of the volumes of the RGVPs reconstructed with T1w-based RGVP segmentation (Section 2.3.3). In addition, to assess the reliability of the ROI selection, we performed an inter-expert reliability validation (Section 2.3.4).

#### 2.3.1 Reconstruction rate of RGVP subdivisions

In this experiment, we assessed the ability of each tractography method to successfully reconstruct the four subdivisions of the RGVP. Specifically, for each RGVP subdivision (LL, RR, LR and RL), we considered that the fiber pathway was successfully traced if there was at least one fiber retained after ROI selection. Then, across all 57 subjects under study, the percentage of successfully traced pathways for each subdivision was computed as the reconstruction rate. In addition to the reconstruction rate per subdivision, we also computed the overall reconstruction rate for the full RGVP pathway (i.e., the percentage of subjects where all four RGVP subdivisions were successfully reconstructed). A statistical comparison was performed across the four tractography methods in terms of the overall reconstruction rate, as follows. First, the overall number of detected subdivisions was measured for each subject and tractography method. (An overall number of 0 indicated no subdivisions were detected, and 4 indicated that all subdivisions were found.). After that, a one-way repeated measures analysis of variance (ANOVA) was performed across the four methods. Following that, two-group Cochran’s Q tests (Cochran, 1950) were performed between each pair of the four tractography methods (a total of 6 comparisons), with a false discovery rate (FDR) correction across the 6 comparisons. (Cochran’s Q test is a non-parametric statistical comparison method for differences between two or more groups of matched samples, where each sample has a binary outcome, e.g. 0 and 1.).

#### 2.3.2. Percentage of decussating fibers

In this experiment, we computed the percentage of the decussating and non-decussating fibers to compare the performance of the four tractography methods in the optic chiasm. It has been suggested that in neurotypical cases there are more decussating fibers than non-decussating fibers (approximately 53% to 58% of the RGVP fibers decussate at the optic chiasm (Chacko, 1948; Kupfer et al., 1967; v. Sántha, 1932)). As a result, multiple studies have investigated quantification of fiber decussation in the optic chiasm using tractography (Ather et al., 2019; Puzniak et al., 2019; Roebroeck et al., 2008; Staempfli et al., 2007). In our study, for each tractography method, we computed the percentage of decussating fibers per subject. A statistical comparison was performed across the four tractography methods, as follows. A one-way repeated measures analysis of variance (ANOVA) was computed across the four methods to determine whether they were statistically different from each other. Following that, we performed a post-hoc paired t-test between each pair of the four tractography methods (a total of 6 comparisons), with FDR correction across the 6 comparisons.

#### 2.3.3 Correlation between T1w-based and tractography-based RGVP volumes

Anatomical T1w images have been widely used to estimate the volume of the RGVP (Simon J. Hickman et al., 2004; Kolbe et al., 2009; Ramli et al., 2014). In this experiment, we assessed the correlation of the volume of the RGVP reconstructed using tractography with T1w-based RGVP segmentation. For computing the volume of the T1w-based segmentation, first an RGVP segmentation was created on a T1w template image (MNI152) by a clinical expert (S.Y.), as shown in Figure 5. This RGVP segmentation was then applied to each individual subject using a registration between the T1w template and the subject-specific T1w data using ANTs (Avants et al., 2009). From the registered T1w-based segmentation, we computed the T1w-based RGVP volume. For computing the volume of the RGVP reconstructed using tractography, we created a normalized voxel-wise tract density map, where the value represents the number of fibers passing through each voxel divided by the maximum number of fibers across all voxels. This tract density map was converted into a binary map using a range of thresholds (from 0.05 to 0.25, using an increment of 0.05) to eliminate influence of voxels passed by only a few fibers (these are usually false positive fibers, and this procedure is widely used in other studies (Reid et al., 2020; Tian et al., 2018)). From this binary map, we computed the tractography-based RGVP volume. Then, for each tractography method, we computed Pearson’s correlation between T1w-based and tractography-based RGVP volumes across all subjects. We also computed the mean absolute error (MAE) between the T1w-based and tractography-based volumes to evaluate their absolute differences. The correlation coefficient and the p-value are reported for each method. (To evaluate the effect of the registration, we also repeated the experiment using an additional widely used registration algorithm provided in FSL (Jenkinson et al., 2012) (see Supplementary Materials 4 and 5). Our results showed using ANTs generated a better result where more significant correlations were obtained; thus, in the paper, we report the results obtained using ANTs. However, we note that our overall results regarding the RGVP volume correlations as well as the Normalized Overlap Score (NOS) experiment introduced below (Section 2.3.6) are not particularly sensitive to the choice of registration algorithm employed. See Supplementary Materials 4 and 5 for detailed results and discussion.)

**Figure 5.**
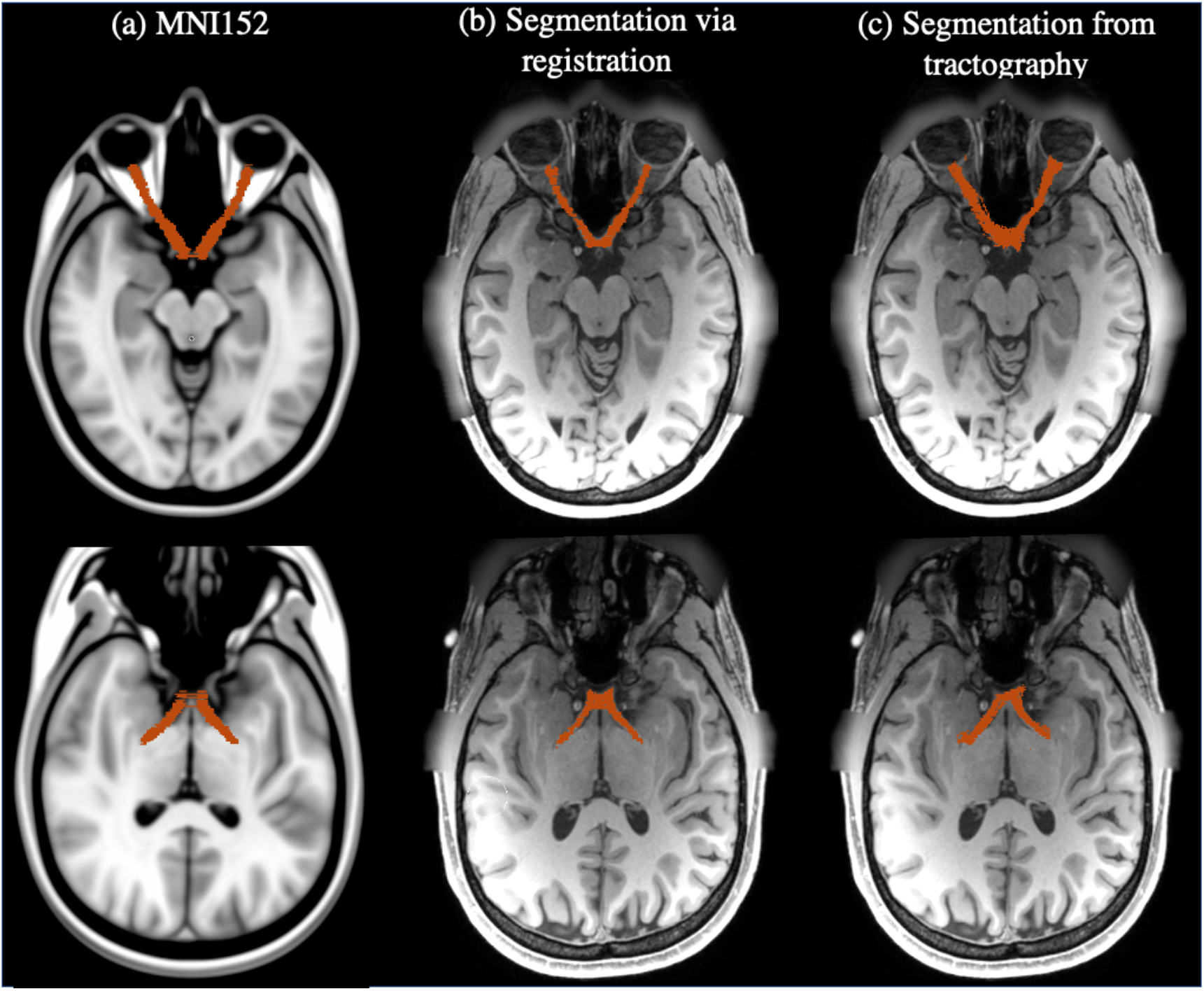
Visualization of RGVP segmentations. (a) shows the expert RGVP segmentation in MNI space; (b) shows a subject’s RGVP segmentation, which was warped from the RGVP segmentation in MNI space. (c) shows a RGVP segmentation computed from tractography results.

### 2.3.4 Inter-expert validation

To assess the reliability of the RGVPs reconstructed for each tractography method, we conducted an inter-expert reliability validation. This experiment evaluated the spatial overlap of the RGVPs selected by two different experts, which is a common strategy in previous studies of tractography (Benjamin et al., 2014; Davis et al., 2009; Dayan et al., 2015; Lilja et al., 2014; Voineskos et al., 2009; Xie et al., 2020b). For each subject under study, a new set of ROIs was drawn by an additional expert (S.Y.) and used for selecting the second RGVP (as described in Section 2.2.2). Then, for the two RGVPs from the two experts, we computed the weighted Dice (wDice) score (Cousineau et al., 2017a) to measure their spatial overlap. wDice is a metric designed specifically for measuring tract spatial overlap (Cousineau et al., 2017a; Zhang, Cetin Karayumak, et al., 2020; Zhang, Wu, et al., 2019). wDice score extends the standard Dice score (Dice, 1945) taking account of the number of fibers per voxel so that it gives higher weighting to voxels with dense fibers, as follows:

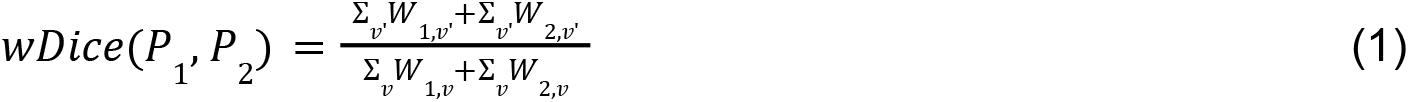

Where *P*_1_, *P*_2_ represent two RGVPs selected using different expert-drawn ROIs, *v’* indicates the set of voxels that are within the intersection of the volumes of *P*_1_ and *P*_2_, *v* indicates the set of voxels that are within the union of the volumes of *P*_1_ and *P*_2_, and *W* is the fraction of the fibers passing through a voxel. A high wDice score represents a high reproducibility between the two RGVPs identified. A statistical comparison was performed across the four tractography methods, as follows. A one-way repeated measures analysis of variance (ANOVA) was computed across the four methods to determine whether they were statistically different from each other. Following that, we performed a post-hoc paired t-test between each pair of the four tractography methods (a total of 6 comparisons), with FDR correction across the 6 comparisons.

### 2.3.5 Expert judgment

We performed an expert judgment experiment to evaluate the quality of the RGVP reconstructions obtained by the four tractography methods, following the expert tract ranking method in our previous work (Gong et al., 2018). One expert (S.Y.) visually ranked the four RGVPs from each subject, as follows. The four tractography files were loaded into the 3D Slicer software (Kikinis et al., 2014; Norton et al., 2017; Zhang, Noh, et al., 2020), overlaid on the anatomical T1w and b0 images. The expert was blinded to the origin of each RGVP: tractography files were anonymized so that there was no information about which RGVP was from which tractography method. The overall RGVP reconstruction quality ranking was performed according to: 1) whether the reconstructed RGVP was anatomically reasonable as appearing on the T1w image, 2) whether there were apparent outlier fibers, and 3) whether the fiber density was sufficient to cover the RGVP pathway. The expert was asked to rank the four reconstructions based on his judgment, where a rank of 1 was the best and 4 was the worst. If two reconstructions were equally good or bad to the expert, these two RGVP reconstructions were given the same rank score. To summarize the expert judgment results, for each method, we then computed the mean and the standard deviation of the ranking scores.

### 2.3.6 Normalized Overlap Score to Quantify Tractography Spatial Agreement

To quantify the RGVP reconstruction performance at the group level, we conduct an experiment using the normalized overlap score (NOS) (D. Q. Chen, Zhong, et al., 2016). NOS is a quantitative metric that measures the spatial agreement of tractography across multiple subjects, and it has been demonstrated to be a stable rating metric to assess the overlap of cranial nerve tractography across subjects. In our experiment, for each tractography method, the reconstructed RGVP of each individual subject was converted to a binary volumetric image, where the value 1 represents there is at least one fiber streamline passing through the voxel. The converted binary images of all subjects under study were co-registered to the MNI template space by registering each subject’s baseline image to the MNI152 template using ANTs (Avants et al., 2009). These binary images were stacked together to form a conjunction image. Each voxel value *s* in the conjunction image denotes the percentage of subjects that have RGVP fibers passing through the voxel, representing the RGVP reconstruction overlap value across subjects (range from 0 to 100% of subjects). The NOS was then calculated by thresholding the conjunction image at multiple thresholds, as follows:

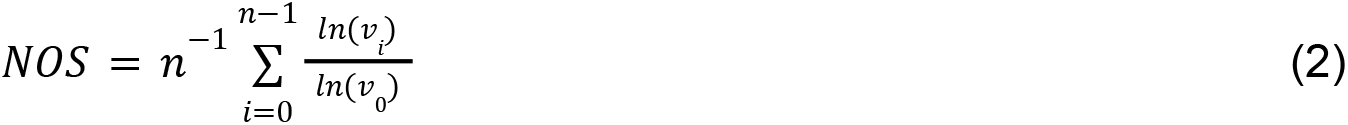

where *n* is the number of thresholds (n=10 is used as suggested in (D. Q. Chen, Zhong, et al., 2016)), *v*_i_ is the number of voxels with overlap value *s* higher than the *i*-th overlap threshold (i.e.,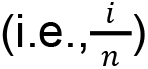).

## 3. Experimental results

### 3.1 Reconstruction rate of RGVP subdivisions

Figure 6 shows the reconstruction rate for each of the four RGVP subdivisions, as well as the overall reconstruction rate across all four subdivisions and all subjects. iFOD1 generated the highest overall reconstruction rate (94.74%), followed by UKF-2T (92.98%). This was followed by SD-Stream (70.18%), and UKF-1T (50.88%). The ANOVA comparison showed that the overall reconstruction rates were significantly different across the four methods. Post-hoc paired t-tests (with FDR correction) showed that the reconstruction rates of both UKF-2T and iFOD1 were significantly higher than UKF-1T and SD-Stream, while there were no significant differences between UKF-2T and iFOD1. The reconstruction rates of each RGVP subdivision showed the same pattern as the overall reconstruction rates (i.e., iFOD1 was the highest, followed by UKF-2T, SD-Stream, and then UKF-1T). In the UKF-2T and iFOD1 methods, the reconstruction rates across the four subdivisions are similar, while in the UKF-1T and the SD-Stream methods, the reconstruction rates of the RL and LR subdivisions (decussating fibers) are on average lower than those of the LL and RR subdivisions (non-decussating fibers).

**Figure 6.**
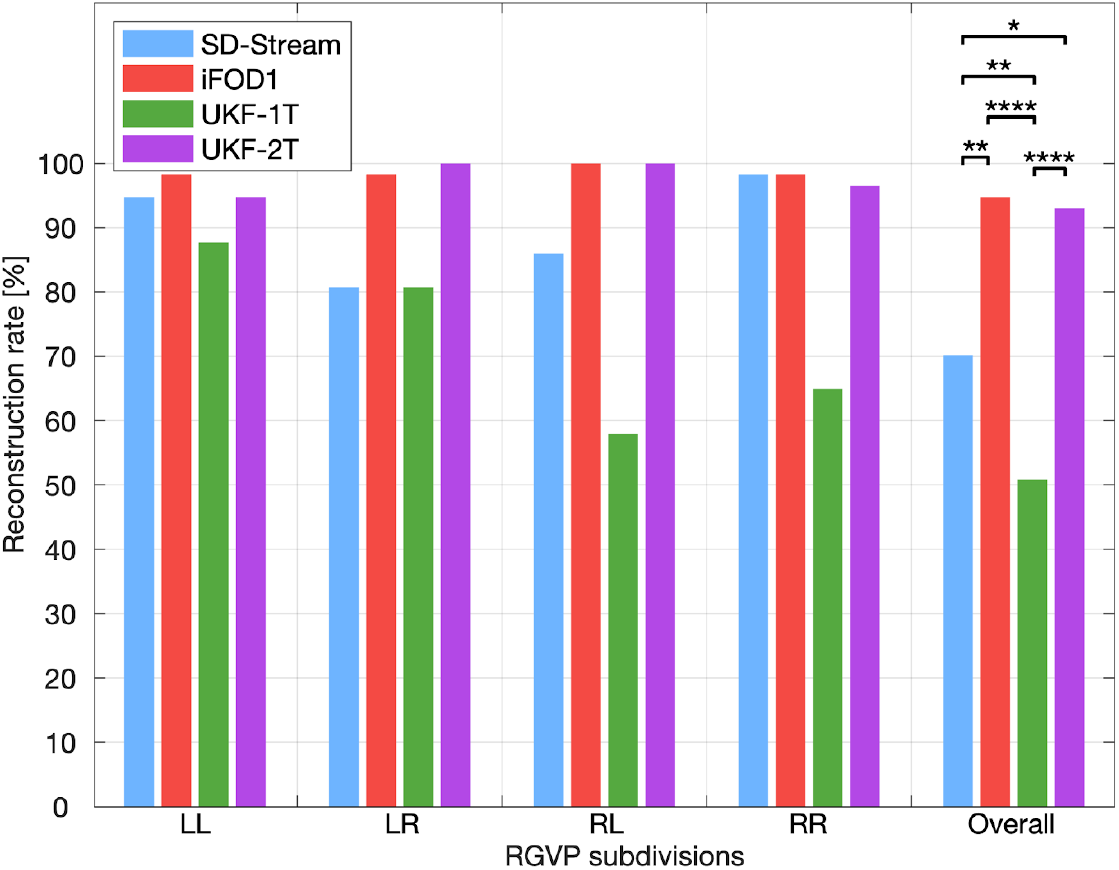
Reconstruction rate of RGVP subdivisions using different tractography methods. The overall RGVP reconstruction rates were statistically significantly different across the four compared tractography methods (ANOVA, p<0.0001). Post-hoc two-group Cochran’s Q tests (with FDR correction) with significant results are indicated by asterisks. *: p<0.05; **: p<0.01; ****: p<0.0001.

### 3.2 Percentage of decussating fibers

Figure 7 gives the results for the percentage of decussating fibers obtained by each tractography method. The decussating percentages of UKF-2T and iFOD1 are higher than 50%, showing more decussating fibers than non-decussating fibers (corresponding to the RGVP anatomy as reported in previous studies (Cajal, 1899; Chacko, 1948; Kupfer et al., 1967)), while the decussating percentages of SD-Stream and UKF-1T are lower than 50%.

**Figure 7.**
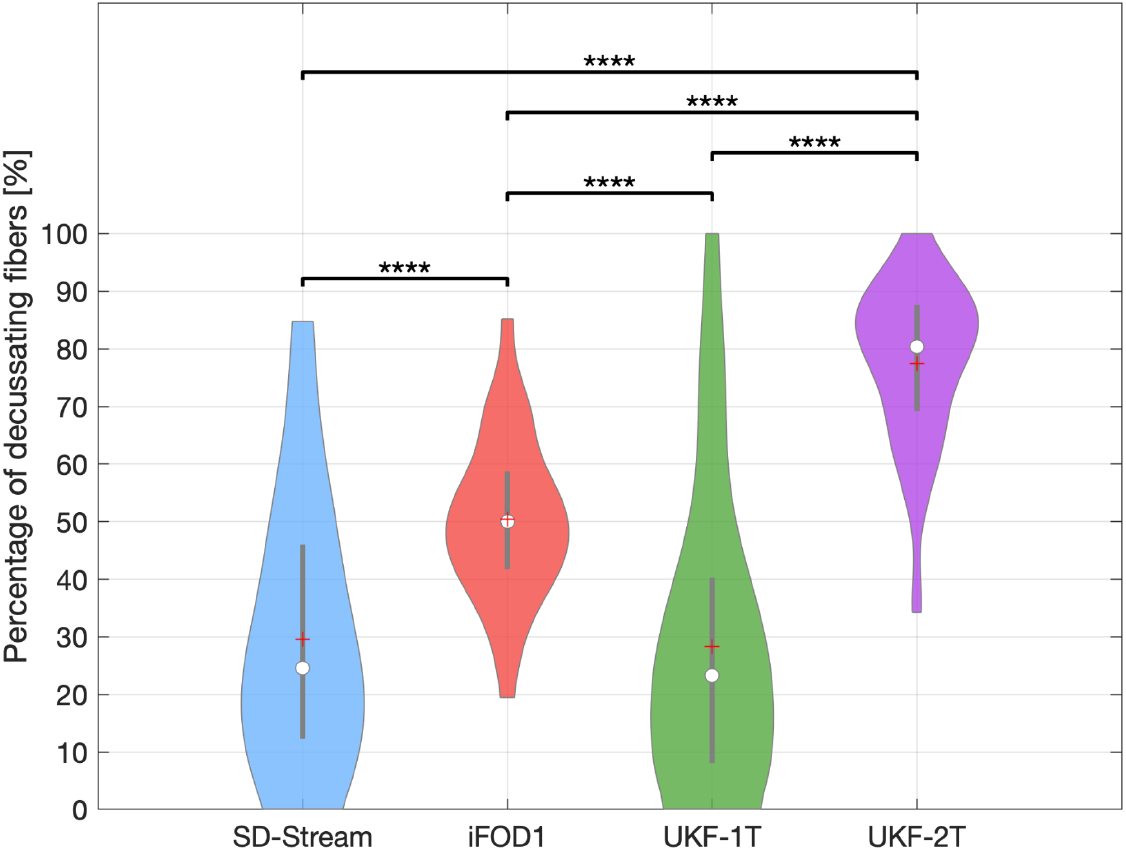
Boxplot indicates the percentage of decussating fibers of all subjects using each tractography method. On each box, the central line indicates the median, the plus symbol indicates the mean, and the bottom and top edges of the box indicate the 25th and 75th percentiles, respectively. The percentage of decussating fibers was statistically significantly different between the different tractography methods (ANOVA, p<0.0001). The Post-hoc paired t-tests (with FDR correction) with significant results are indicated by asterisks. **: p<0.01; ****: p<0.0001.

### 3.3 Correlation between T1w-based volume and tractography-based volume

Figure 8 gives the results of the correlation analysis and the MAE results between the T1w-based volume and the tractography-based volume of the RGVP. No significant correlations were obtained by the UKF-1T or iFOD1 methods at any of the threshold values. The SD-Stream and UKF-2T methods obtained significant correlations. The UKF-2T method obtained significant correlations at all five threshold values, whereas the SD-Stream method obtained significant correlations at four of the threshold values. In addition, the UKF-2T method obtained a lower MAE at each of the threshold values compared to the SD-Stream method. Overall, these results show that, across the four compared methods, the UKF-2T method generated the tractography-based volume that was most similar to the T1w-based volume.

**Figure 8.**
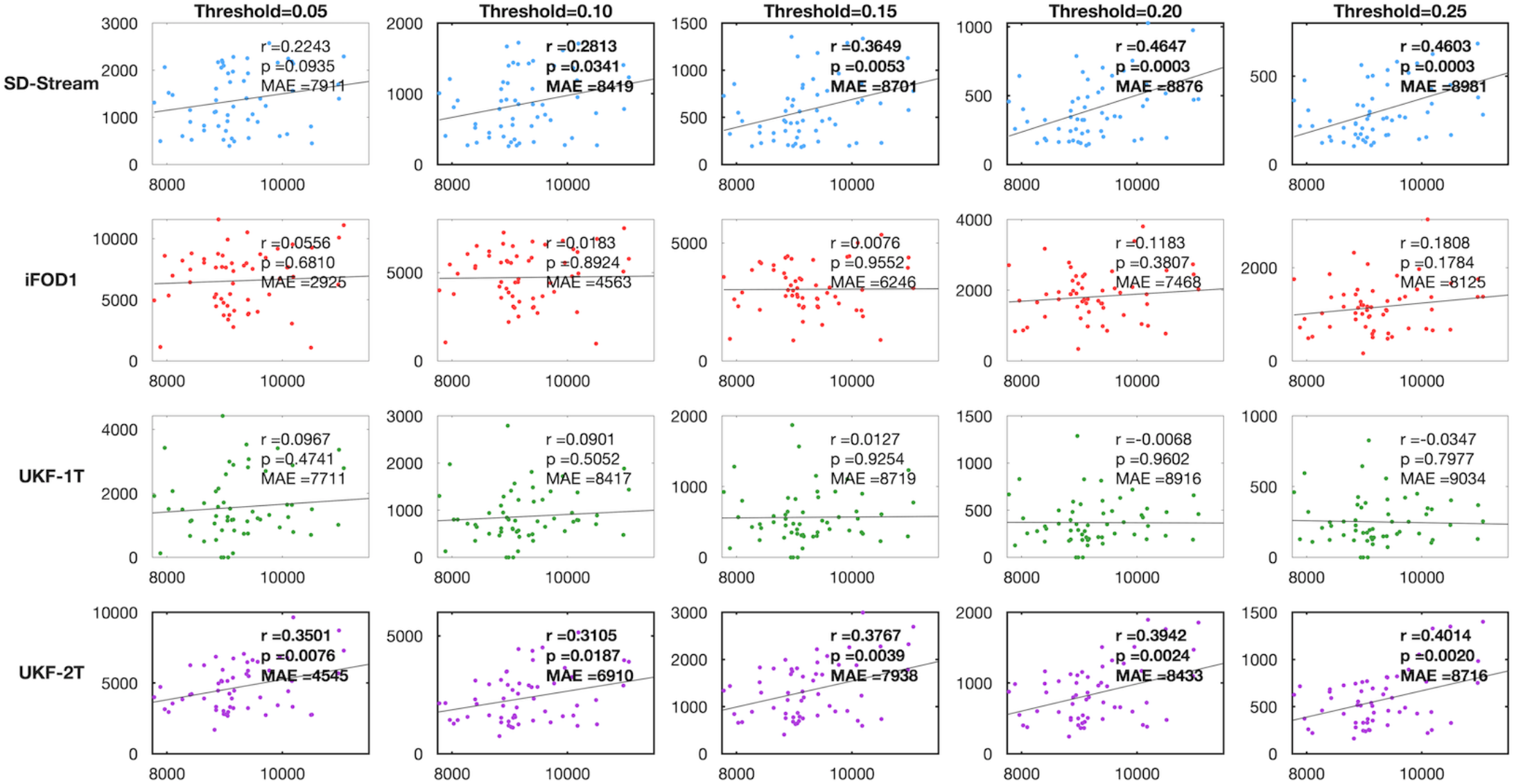
Scatter plots of correlation between T1w-based volume and tractography-based volume. In each plot, the y-axis shows the tractography-based volume (mm^3^) and the x-axis shows the T1w-based volume (mm^3^). Each row represents the correlation results of one tractography method under different threshold values. The correlation coefficient r and the p-value are reported for each plot. Plots showing significant correlations are outlined in bold. MAE (mean absolute error) between the tractography-based and the T1w-based volumes across all subjects is also reported.

### 3.4 RGVP visualization

Figure 9 gives a visual comparison of the RGVP results using different tractography methods. In general, the CSD-based tractography methods (SD-Stream and iFOD1) generated a large number of visually apparent false positive fibers, while the UKF tractography methods (UKF-1T and UKF-2T) generated more anatomically plausible RGVP fibers, corresponding to the RGVP as appearing on the T1w image.

**Figure 9.**
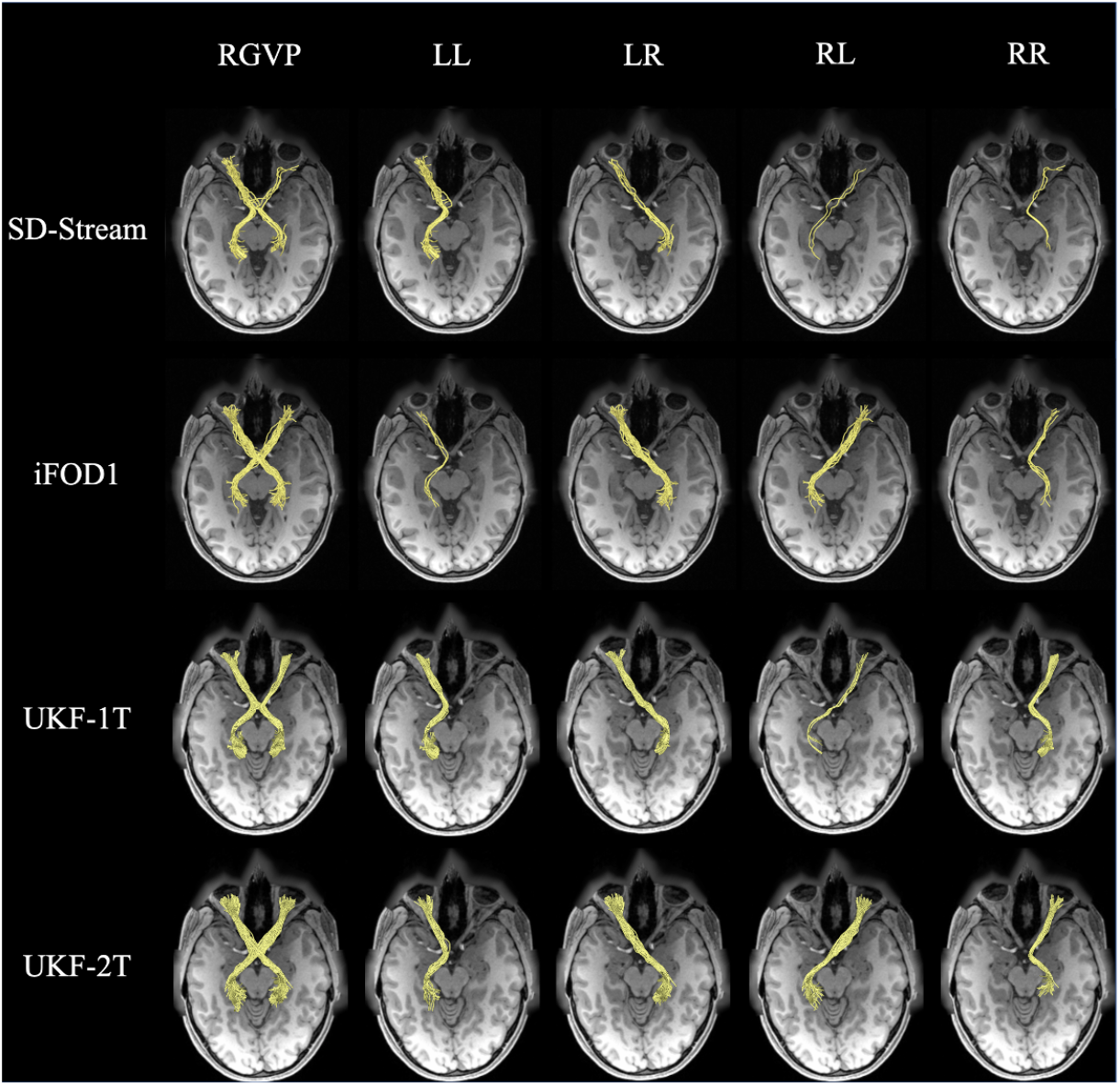
Visual comparison of the RGVP reconstructed using the four tractography methods. The RGVPs (yellow bundles) obtained from one HCP subject are displayed, overlaid on the T1w image. Each row shows the RGVP and its subdivisions using one of the tracking strategies. The first column shows the overall RGVP fiber pathway, and the following columns show the four subdivisions.

### 3.5 Inter-expert validation

Figure 10 gives the inter-expert validation results. The wDice scores of the two UKF tractography methods were the highest (over 0.81), followed by the SD-Stream method (0.75), while the iFOD1 method obtained the lowest wDice score (0.74) (wDice = 0.72 has been suggested to be a threshold for a good tract spatial overlap (Cousineau et al., 2017a)). We note that an ANOVA analysis shows that there are no significant differences across the four methods, indicating that all tractography methods show good tract spatial overlap in inter-expert validation.

**Figure 10.**
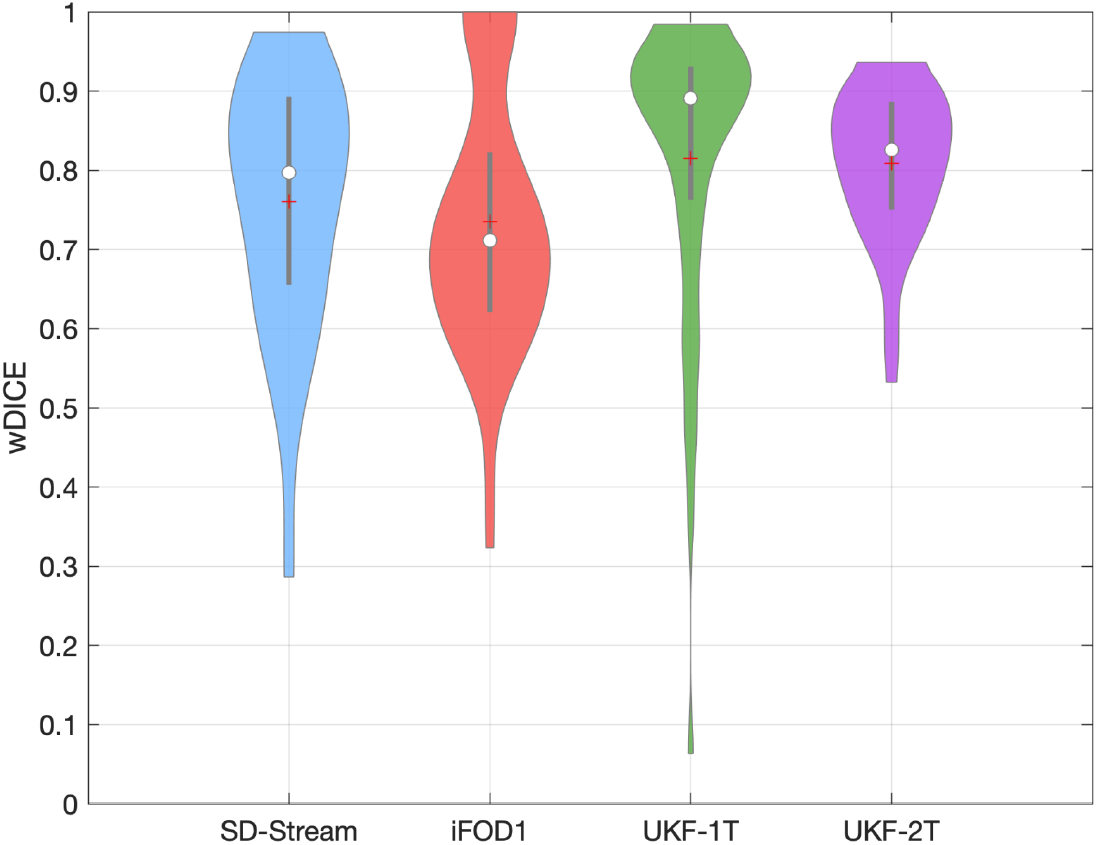
Violin plot indicates inter-expert validation results. The wDice score was statistically significantly different between the different tractography methods (ANOVA, p=0.06).

### 3.6 Expert judgment

Expert judgment was performed to rank the RGVP reconstructions of the 57 subjects under study. The rank could range from 1 to 4, where 1 was the best-judged reconstruction and 4 was the worst-judged reconstruction. The averaged ranking scores of the SD-Stream, iFOD1, UKF-1T and UKF-2T methods were 3.26±0.72, 2.93±0.70, 2.32±1.38, and 1.47±0.50, respectively (shown in Figure 11). An ANOVA comparison indicates that the overall reconstruction rates are significantly different across the four tractography methods. Post-hoc paired Wilcoxon signed-rank tests (with FDR correction) show that UKF-2T has a significantly higher expert judgment score than the other three methods. These results indicate that the UKF-2T method obtained the best expert evaluation performance in this experiment.

**Figure 11.**
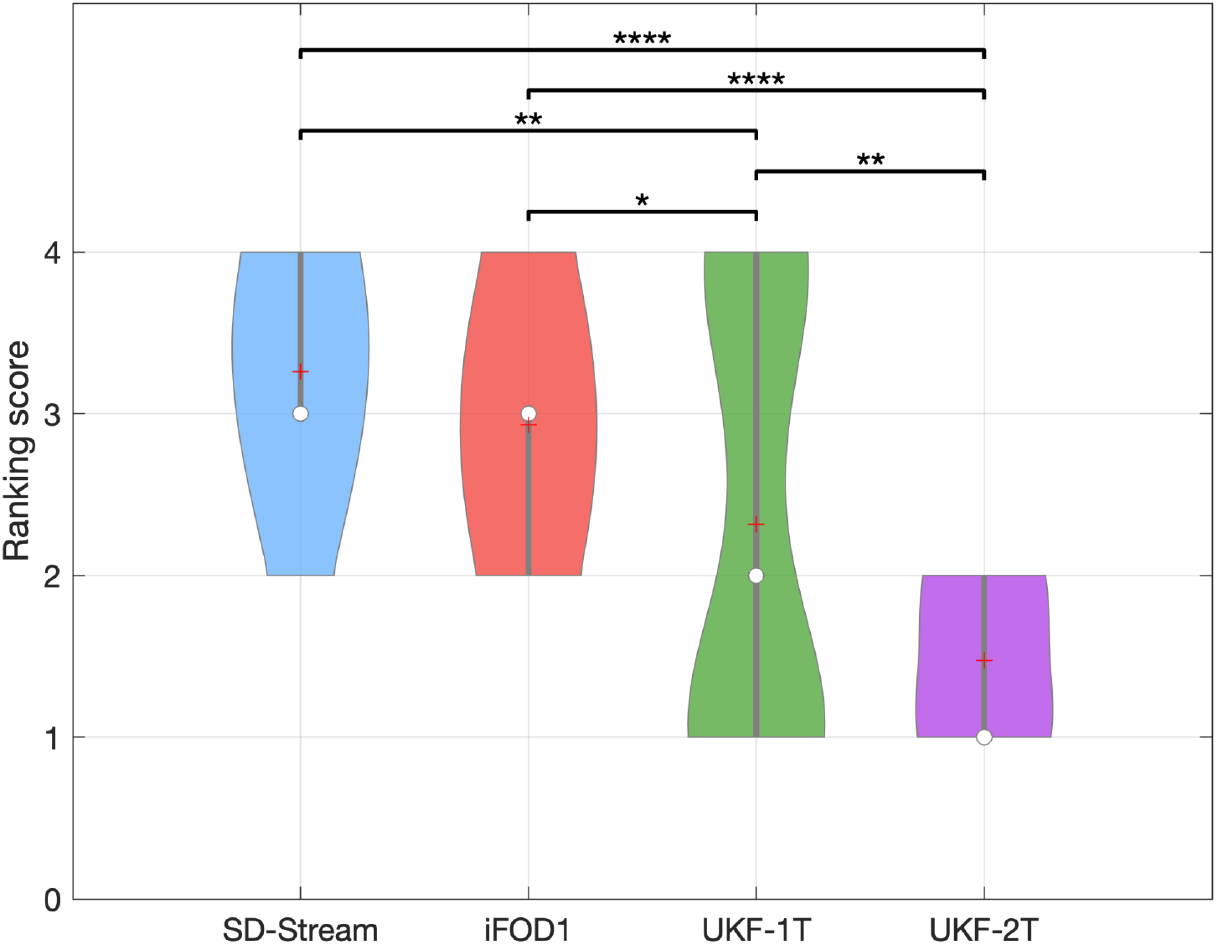
Violin plot indicates the ranking score of all subjects using each tractography method. On each violin box, the central circle indicates the median, the plus symbol indicates the mean, and the bottom and top of the gray central vertical line indicate the 25th and 75th percentiles, respectively. The ranking score was statistically significantly different between the different tractography methods using the Friedman test (p<0.0001). Post-hoc paired Wilcoxon signed-rank tests (with FDR correction) with significant results are indicated by asterisks. *: p<0.05; **: p<0.01; ****: p<0.0001.

### 3.7 Normalized overlap score of each tractography method

Figure 12 gives the results of the NOS comparison across different tractography methods. In each subfigure, the log ratio curve is provided, from which the area under the curve is computed, i.e. the NOS. The UKF-2T method obtained the highest score (NOS=0.718), indicating the highest overlap of tractography across subjects. The next highest scores were obtained by iFOD1 (NOS=0.605) and UKF-1T (NOS=0.508). SD-Stream obtained the lowest score (NOS=0.398).

**Figure 12.**
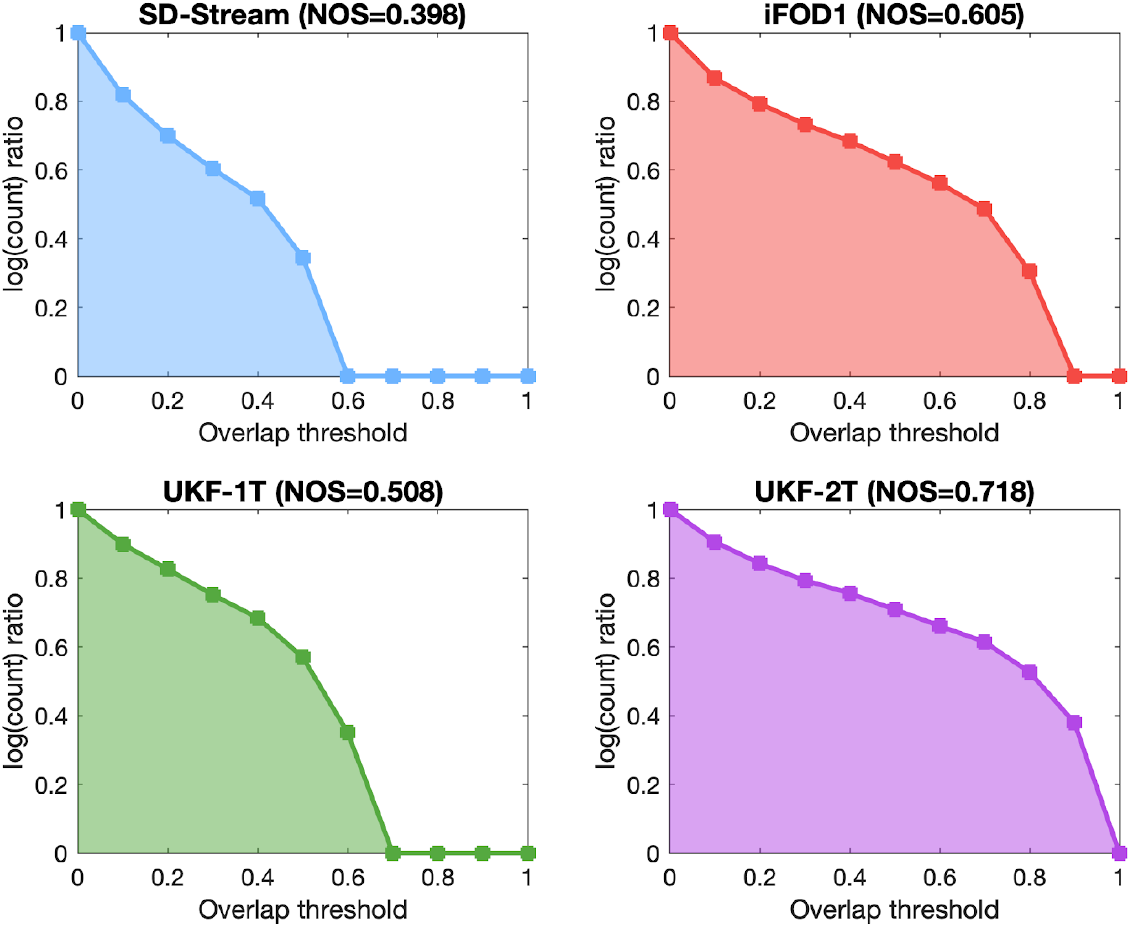
NOS of conjunction images generated by different tractography methods. The y-axis shows the log ratio (the ratio that is summed in equation (2)), while the x-axis shows different threshold values of conjunction images. The NOS is the area under the curve.

## 4. Discussion

In this study, we proposed, for the first time, to investigate the performance of multiple tractography methods for reconstruction of the complete RGVP including the four anatomical subdivisions. We compared the RGVP reconstructions of four tractography methods, including deterministic (SD-Stream) and probabilistic (iFOD1) tractography based on the CSD model, and UKF tractography with one-tensor (UKF-1T) and two-tensor (UKF-2T) models. Anatomical measurements were conducted to assess the advantages and limitations of the four tractography methods for reconstructing RGVP. To this end, we computed the reconstruction rate of the four major RGVP subdivisions, the percentage of decussating fibers at the optic chiasm, and the correlation between the volumes of traced RGVPs and T1w-based RGVP segmentations. Finally, we performed an expert judgment experiment to rank the anatomical appearance of the reconstructed RGVPs, and we quantified the performance of each tractography method across subjects using the normalized overlap score method. Overall, we have three main observations. First, across the four compared methods, we found that UKF-2T and iFOD1 obtained better performance than UKF-1T and SD-Stream. The RGVP reconstruction rates of UKF-2T and iFOD1 (over 92% in both methods) were significantly higher than UKF-1T and SD-Stream (50.88% and 70.18%, respectively). UKF-2T and iFOD1 also traced more decussating than non-decussating fibers in the optic chiasm, which is consistent with the findings of previous studies (approximately 53% to 58% as reported in (Chacko, 1948; Kupfer et al., 1967; v. Sántha, 1932)), whereas UKF-1T and SD-Stream trace more non-decussating fibers than decussating fibers. Second, comparing between UKF-2T and iFOD1, the percentage of decussating fibers in the iFOD1 method was more similar to the reported range in the previous studies, while the UKF-2T method obtained better volume correlation with the T1w-based RGVP segmentation. Third, the UKF-2T method obtained visually better RGVP pathways as appearing on the T1w image and was the top-performing method according to the normalized overlap score. Below, we discuss several detailed observations regarding the comparison results for the four tractography methods.

We demonstrated that multi-fiber tractography (UKF-2T) had advantages for reconstructing the complete RGVP compared to the single-fiber model (UKF-1T) within the UKF tractography framework. This is likely because a single-fiber model cannot estimate multiple fiber orientations (a well-known issue in the tractography literature (A. L. Alexander et al., 2001; D. C. Alexander, 2005; Behrens et al., 2007; Jeurissen et al., 2019; J.-D. Tournier et al., 2004; Tuch et al., 2002)) in the region of the optic chiasm, where the decussating fibers cross. Multiple studies have suggested that multi-fiber models are beneficial for cranial nerve reconstruction (Behan et al., 2017; D. Q. Chen, DeSouza, et al., 2016; Z. Chen et al., 2016; Xie et al., 2020a). However, multi-fiber tractography is expected to produce more outliers than single-fiber tractography (Maier-Hein et al., 2017). This can be seen when comparing UKF-2T vs UKF-1T in Figure 9. Due to the presence of outlier fibers, UKF-2T also has a lower wDice score than UKF-1T (Figure 10).

We showed that probabilistic tractography (iFOD1) was more effective for reconstructing the RGVP than deterministic tractography (SD-Stream) within the CSD tractography framework. This is potentially because the deterministic tractography could only follow the principal orientation along the FOD amplitude peak, thus it could not effectively track through the skull base region where the signal-to-noise ratio (SNR) is low. On the other hand, the probabilistic tractography allowed fiber tracking along multiple orientations for each FOD. Previous tractography-based cranial nerve studies have also shown the advantages of probabilistic tractography over traditional deterministic tractography, in particular, due to its good performance to handle the low SNR in the skull base (Jacquesson, Cotton, et al., 2019; Rueckriegel et al., 2016; Zolal et al., 2017).

Comparing between the UKF-2T and iFOD1 methods (the two methods that in general obtained the best RGVP reconstruction performance), the UKF-2T method produces reconstructed RGVPs that were judged to better correspond to known anatomy. In UKF-2T, each tracking step employs prior information from the previous step to help stabilize model fitting and includes a probabilistic prior about the rate of change of fiber orientation (defined as the parameter *Qm* introduced above). Consequently, sharp fiber curvatures are avoided as they are very unlikely, whereas fiber curvatures (e.g., curvature at the optic chiasm) supported by the dMRI are still allowed. This tracking strategy can reduce the effect of noise, susceptibility artifacts, and partial volume averaging along the RGVP between the optic chiasm and the optic nerve (Bender et al., 2014; Yoshino et al., 2016), where it is surrounded by bone and air (see Supplementary Material 3). Thus, UKF-2T generated highly smooth fiber streamlines that are the anatomically expected shape of the RGVP fibers as appearing on the T1w images. However, using prior information from the previous step can encourage straight fibers and therefore tend to produce more decussating RGVP fibers. Our study showed that using UKF-2T reconstruction, the percentage of decussating fibers was higher than that known from anatomical dissection studies. (A similar tracking strategy is used in the iFOD2 method; in our initial experiment, the non-decussating fibers were often not traced, resulting in incomplete detection of RGVP subjects in over half of the subjects (Supplementary Material 1).). The iFOD1 method performs tractography following randomly sampled directions. As a result, the percentage of decussating fibers was unbiased, and the result was close to that known from studies of anatomy (Chacko, 1948; Kupfer et al., 1967; v. Sántha, 1932). However, the traced fiber streamlines were highly curved, where certain fiber segments were outside the RGVP pathway (see Figure 9). Overall, these results indicate that it is challenging to accurately track RGVP fibers that correspond to known anatomy and have an approximately correct percentage of decussating fibers. Researchers and clinicians may benefit from tailoring tractography algorithm selection depending on the target applications. The UKF-2T method could be more useful in qualitative applications, e.g., pre-surgical planning in the setting of skull base tumors affecting the RGVP (Lober et al., 2012; Potgieser et al., 2014), as it can better reconstruct the known anatomy of the RGVP. The iFOD1 method could be more useful in quantitative studies that assess RGVP decussation abnormalities in brain diseases (e.g., albinism (Puzniak et al. 2019)), as it can better estimate the percentage of decussating fibers.

In this work we used the popular expert selection strategy to identify the RGVP based on multiple expert-identified ROIs. These were separately located in the optic nerve, optic chiasm, and optic tract. Note that the ROIs in the optic tract are different from many previous studies, which defined the LGN by performing nonlinear registration of the Juelich histological atlas (Haykal et al., 2019; Maleki et al., 2012; Millington et al., 2014; Staempfli et al., 2007). We suggest that using the LGN as an ROI may not be a good choice for RGVP selection, whether it is generated automatically or manually. On the one hand, registration bias and fuzziness of a probabilistic atlas may increase the inaccuracy of the LGN boundary. On the other hand, it is challenging to accurately recognize the LGN on current medical image data. Using the LGN as an ROI for RGVP selection may therefore generate more false positive fibers, especially when selecting fibers generated by tractography using multi-fiber models. Several studies have traced the RGVP in clinical data with lower resolution than the HCP data employed here; in these studies, various approaches for fiber cleaning were necessary after fiber selection (Allen et al., 2018; Malania et al., 2017; H. Takemura et al., 2019).

Our inter-expert validation showed overall a high reliability of the RGVP reconstruction results. In particular, the wDice scores of all tractography methods were over 0.72, which is a threshold for good tract spatial overlap (Cousineau et al., 2017b).

In this work, we performed what we believe is the first study to investigate the performance of multiple tractography methods for reconstruction of the complete RGVP including the four anatomical subdivisions. In the literature of tractography-based RGVP studies, most studies have focused on certain RGVP subregions of interest, e.g., the optic nerve, the optic tract, and/or the RGVP fibers within the optic chiasm (as summarized in Table 1). Several studies have shown successful tracking of the complete RGVP fiber pathway including the four major anatomical subdivisions (i.e., the two decussating and the two non-decussating fiber pathways) (Altintaş et al., 2017; Hofer et al., 2010; Lober et al., 2012; Panesar et al., 2019; Yoshino et al., 2016). Some of these studies demonstrated that RGVP reconstruction could provide valuable information for pre-surgical planning. Nevertheless, these studies mostly present a visual demonstration, e.g., a visualization of RGVP fiber displacement due to tumors (Lober et al., 2012; Potgieser et al., 2014), without a comprehensive evaluation of different tractography methods. In this study, the higher reconstruction rate and high performance according to expert judgment indicated that UKF-2T is potentially a better tractography method for pre-surgical planning. Other previous studies (Ather et al., 2019; Puzniak et al., 2019) have reported the relationship between albinism and RGVP reconstruction results. However, the percentage of decussating fibers of these studies is underestimated compared to that known from histological studies (Chacko, 1948; Kupfer et al., 1967; v. Sántha, 1932). Our study showed that using iFOD1 reconstruction, the percentage of decussating fibers was close to the known anatomy (Chacko, 1948; Kupfer et al., 1967; v. Sántha, 1932).

Potential limitations of the present study, including suggested future work to address limitations, are as follows. First, in our current study, we focused on the assessment of the four major RGVP subdivisions, while it would be interesting to include more specific RGVP subregions. For example, it would be possible to investigate the RGVP fibers entering the superior colliculus (Purves et al., 2014). Second, we focused on RGVP tracking using dMRI data from healthy adults in the current study. Many research studies have suggested that the RGVP is important for understanding and/or potential treatment of various diseases, including glioma (Hales et al., 2018), optic neuritis (Beck et al., 2003; Brex et al., 2002; Optic Neuritis Study Group, 1997), ischemic optic neuropathy (Attyé et al., 2018; Cho et al., 2016), and sheath meningioma (Schick et al., 2004; Turbin et al., 2002). Further work could include a comparison of different tractography methods on dMRI data from populations with neurological disorders or brain tumors. Third, false positive fiber tracking has been known to be prevalent in tractography (Maier-Hein et al., 2017; Thomas et al., 2014; Xie et al., 2020b). In our study, we have also observed several places where false positive RGVP tracking was likely to happen, in particular, at the RGVP endpoint regions (e.g., fiber tracking into the eyeballs and fiber tracking fanning before entering the LGN). A future investigation could include applying advanced fiber filtering techniques (Daducci et al., 2015; Miller et al., 2019; Pestilli et al., 2014; Smith et al., 2015) to further remove false positive fibers.

## 5. Conclusion

This is the first study to compare multiple tractography methods for reconstruction of the complete RGVP. Overall, we found that the multi-fiber UKF tractography (UKF-2T) and the probabilistic CSD tractography (iFOD1) obtained the best results, showing highly successful performance to identify the four major subdivisions of the RGVP. The iFOD1 method can better quantitatively estimate the percentage of decussating fibers, while the UKF-2T method produces reconstructed RGVPs that are judged to better correspond to known anatomy. However, it is challenging for current tractography methods to both accurately track RGVP fibers that correspond to known anatomy and produce an approximately correct percentage of decussating fibers. We suggest that future tractography algorithm development for RGVP tracking should take consideration of both of these two points.

## Data availability statement

Imaging datasets of 100 subjects from Human Connectome Project (HCP) database (https://www.humanconnectome.org) are used in this paper. The datasets are online available. The computed RGVP tractography data will be made available on request.

## Ethics approval statement

HCP dataset was approved by the ethics committee of Keio University. This study has got the approval of the Ethics Committee to use the HCP data. The certificate stating will provide on request.

## Patient consent statement

No patient datasets are used.

## Permission to reproduce material from other sources

Not applicable.

## Clinical trial registration

Not applicable.

## Acknowledgements

We gratefully acknowledge funding provided by the following National Institutes of Health (NIH) grants: P41 EB015902, P41 EB015898, R01 MH074794, R01 MH111917, R01 MH119222, R01 CA235589, HHSN261200800001E, HHSN26100071, and U01 CA199459; the National Natural Science Foundation of China: 61976190, 61903336; Key Research & Development Project of Zhejiang Province grant: 2020C03070; Major Science and Technology Projects of Wenzhou: ZS2017007; Chinese Postdoctoral Science Foundation (2019M663271). Jianzhong He was supported by a scholarship from the China Scholarship Council (CSC).

