## supplementary for "Comparison of multiple tractography methods for reconstruction of the retinogeniculate visual pathway using diffusion MRI"

**Supplementary Material 1: Comparison of iFOD1 and iFOD2 methods**

The key parameters that may affect the RGVP tractography results were tested in iFOD1 and iFOD2 methods, and a range of values that were large enough to cover possible settings were used for each parameter (shown in Table S1). In this experiment, 20 of 57 HCP subjects were studied. The best-performing parameters, which achieved the highest number of the overall detected RGVP subdivisions, were determined. For iFOD1, the parameter combination: *seedCutoff*=0.006, *stopCutoff*=0.005, and *maximumAngle*=10, achieved the highest number of detected RGVP subdivisions, and for iFOD2 the best parameter combination is: *seedCutoff*=0.006, *stopCutoff*=0.005, and *maximumAngle*=35. The subdivision detection results using these parameter settings are shown in Figure S1. Results indicate significantly better performance of iFOD1 on complete RGVP reconstruction, and therefore iFOD1 is used in the rest of the experiments in this paper.

Table S1. Key parameters and values tested for each tractography method

| **Tractography method** | **Key parameter** | **Description** | **Values tested** |
|---|---|---|---|
| iFOD1 and iFOD2 | seedCutoff | Minimum FOD amplitude for seeding tracking | 0.005 - 0.050 (Increment: 0.005) |
| | stopCutoff | FOD amplitude cutoff for terminating tracking | Equal to or 0.001 lower than the seedCutoff value |
| | maximumAngle | Maximum angle between successive step | 5 - 90 (Increment: 0.005) |

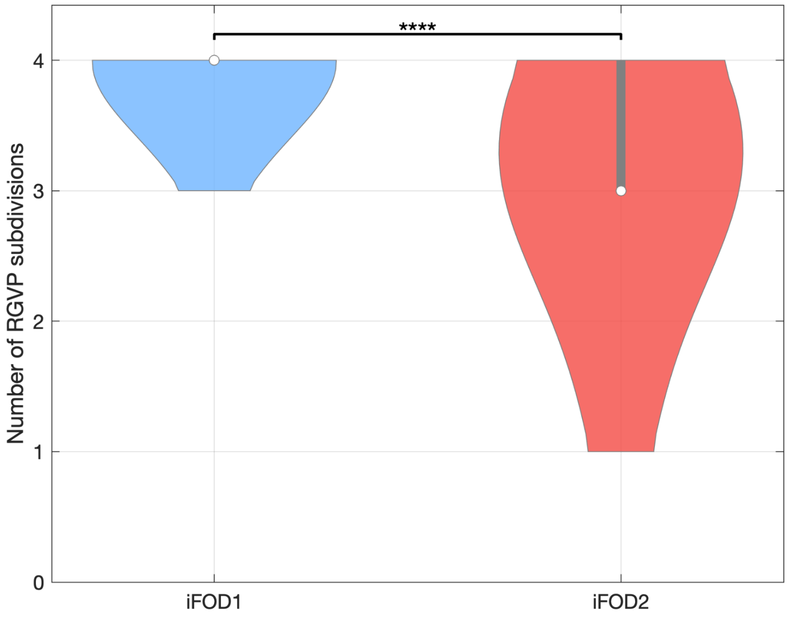


Figure S1. Violin plot indicates the detected number of individual RGVP subdivisions across subjects using iFOD1 and iFOD2 methods, under the best-performing parameter settings for each method. Detected RGVP subdivisions were found to be significantly different between iFOD1 and iFOD2 methods. The asterisk indicates paired t-test across methods. ****: p<0.0001.

**Supplementary Material 2: Experiment to determine best-performing tractography parameters**

Different parameter settings will significantly influence the results of tractography (Parizel et al., 2007). In order to compare multiple tractography algorithms fairly, we first perform an experiment to determine the best-performing parameters for each tractography method, as in our previous work (Xie et al., 2020a).

The key parameters that may affect the RGVP tractography results were tested in each compared method. Detailed information about the parameters is shown in Table S2. For the UKF tractography methods (UKF-1T and UKF-2T), the major parameters include *seedingFA*, *stoppingFA*, *Qm*, and *Ql*. Tractography is seeded in all voxels within RGVP seeding mask (as shown in Figure 3a), where FA is greater than the *seedingFA* threshold value. Tracking stops in voxels where the FA value falls below the *stoppingFA* threshold value. During the tracking, the UKF tractography methods use the *Qm* parameter to control process noise for angles/direction, and use the *Ql* parameter to control process noise for eigenvalues. For the CSD-based methods, the major parameters include *seedCutoff*, *stopCutoff* and *maximumAngle*. Tractography is seeded in the voxels within the mask where the FOD amplitude is higher than the *seedCutoff* threshold value. Tracking stops when the direction of fiber tracking turns through an angle higher than the *maximumAngle* threshold value or the current voxel FOD amplitude falls below the *stopCutoff* threshold value.

In the experiment, in each tractography method, we tested a range of values that were large enough to cover possible settings for each parameter (see Table S2). For each of the 57 subjects (detailed information in Section 2.1) under study, all possible parameter combinations within the selected value ranges were tested. (All other parameters follow the default value suggested by the software.) For each parameter combination, after tractography seeding, we performed ROI-based RGVP fiber selection (as in Section 2.2.2), and we calculated the overall reconstruction rate of RGVP subdivisions (i.e., the percentage of subjects where all four RGVP subdivisions were successfully reconstructed, as in Section 2.3.1). A total of 12,768 putative RGVPs were reconstructed (224 parameter combinations $\times$ 57 subjects) for each of the UKF tractography methods, and a total of 20,520 putative RGVPs were reconstructed in each of the CSD-based tractography methods based tractography method (360 parameter combinations$\times$ 57 subjects) were produced. Then, the best-performing parameter setting was set to those that generated the highest overall reconstruction rate. For each of the SD-Stream and iFOD1 methods, we found two settings that generated the highest overall reconstruction rate, and we chose the setting that is closer to the default setting.

Table S2. Key parameters and values tested for each tractography method

| **Tractography method** | **Key parameter** | **Description** | **Values tested** |
|---|---|---|---|
| SD-Stream and iFOD1 | seedCutoff | Minimum FOD amplitude for seeding tracking | 0.005 - 0.050<br>(increment: 0.005) |
| | stopCutoff | FOD amplitude cutoff for terminating tracking | Equal to or 0.001 lower than the seedCutoff value |
| | maximumAngle | Maximum angle between successive step | 5 - 90<br>(increment: 5) |
| UKF-1T and UKF 2T | seedingFA | Minimum FA for seeding tracking | 0.01 - 0.04<br>(increment: 0.01) |
| | stoppingFA | FA cutoff for terminating tracks | Equal to or 0.01 lower than seedingFA |
| | Qm | Process noise for angles or | 0.001 - 0.004 |

| | | | |
|---|---|---|---|
| | | direction during model fitting | (increment: 0.001) |
| | QI | Process noise for eigenvalues during model fitting | 50 - 350<br>(increment: 50) |

**Supplementary Material 3: The influence of low-SNR dMRI signals on FOD reconstruction**

Here, we show one example of the FOD reconstruction that SD-Stream and iFOD1 methods used for tracking. The dMRI data suffers from noise, susceptibility artifacts and partial volume averaging in the air–bone–tissue interface at the skull base (Bender et al., 2014; Xie et al., 2020b; Yoshino et al., 2016). The FODs shown in Figure S2 were estimated using the CSD model, where we can observe that there are multiple FOD lobes that do not align with the optic nerve pathway (such FOD lobes should be regarded as false-positives).

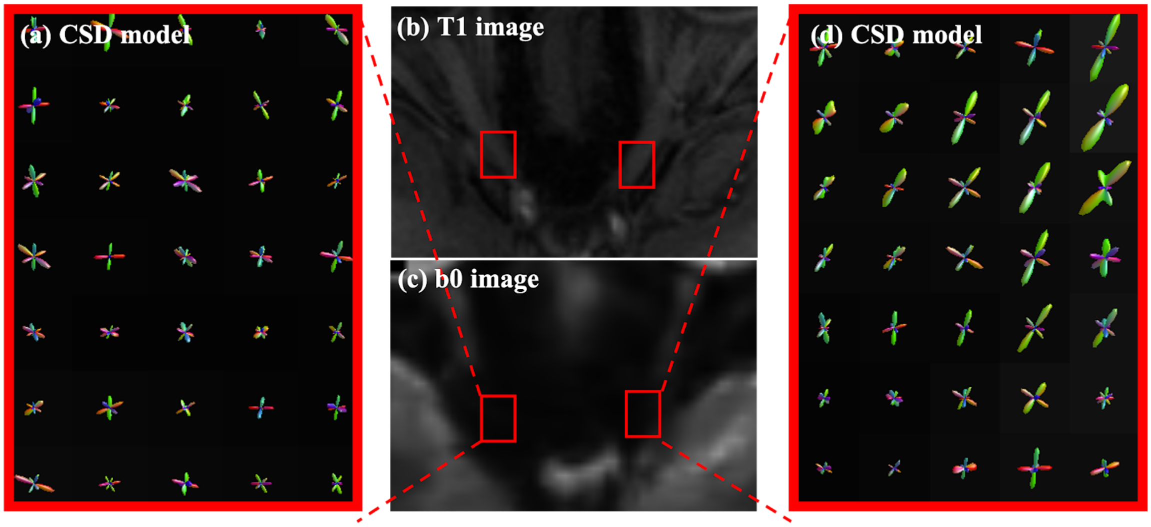


Figure S2. Visualization of FODs at the skull base. (b),(c) Red boxes are located along the RGVP at the skull base; (a), (d) FODs are overlaid on the b0 image.

**Supplementary Material 4: Correlation between T1w-based and tractography-based RGVP volumes (Using FSL)**

In this experiment, we assessed the correlation of the volume of the RGVP reconstructed using tractography with T1w-based RGVP segmentation. For computing the volume of the T1w-based segmentation, first an RGVP segmentation was created on a T1w template image (MNI152) by a clinical expert (S.Y.). This RGVP segmentation was then applied to each individual subject using a registration between the T1w template and the subject-specific T1w data using the non-linear registration algorithm FNIRT/FSL (Andersson et al., 2007; Jenkinson et al., 2012), which uses a b-spline representation of the registration warp field.

Figure S3 gives the results of the correlation analysis and the MAE results between the T1w-based volume and the tractography-based volume of the RGVP. No significant correlations were obtained by the UKF-1T or iFOD1 methods at any of the threshold values. The SD-Stream and UKF-2T methods obtained significant correlations. The UKF-2T method obtained significant correlations at three of the threshold values, whereas the SD-Stream method obtained significant correlations at two of the threshold values. (This result is similar to the results in the main paper using ANTs registration, where UKF-2T showed significant correlations across five threshold values and SD-Stream showed significant correlations across four threshold values).

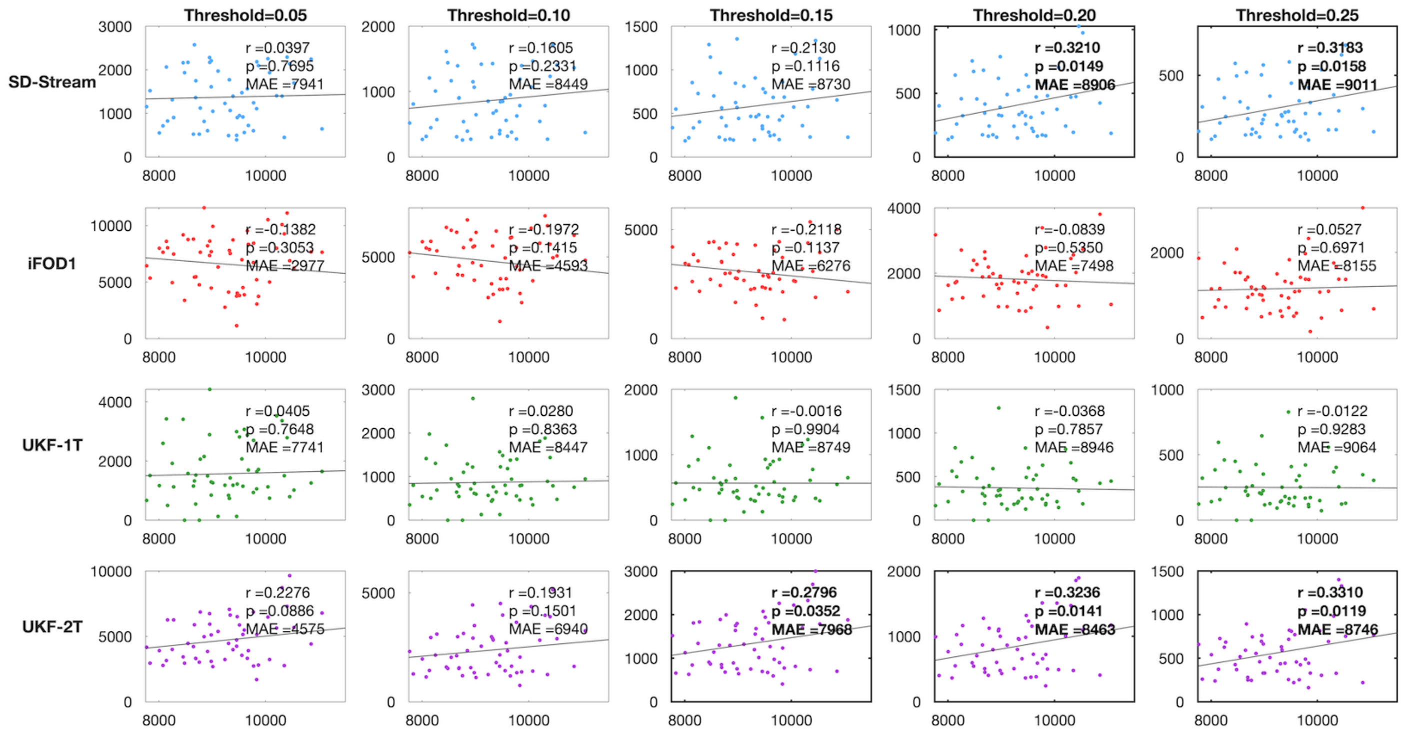


Figure S3. Scatter plots of correlation between T1w-based volume and tractography-based volume. In each plot, the y-axis shows the tractography-based volume (mm$^3$) and the x-axis shows the T1w-based volume (mm$^3$). Each row represents the correlation results of one tractography method under different threshold values. The correlation coefficient r and the p-value are reported for each plot. Plots showing significant correlations are outlined in bold. MAE (mean absolute error) between the tractography-based and the T1w-based volumes across all subjects is also reported.

### Supplementary Material 5: Normalized overlap score (using FSL)

In this experiment, the individual tractography results were mapped to binary spatial images and then transformed to a template space using the non-linear registration algorithm FNIRT/FSL (Andersson et al., 2007; Jenkinson et al., 2012), which uses a b-spline representation of the registration warp field. Conjunction images and the NOSs for each tractography method were computed as in Section 2.3.6. Figure S4 shows the NOS of the SD-Stream, iFOD1, UKF-1T and UKF-2T are 0.268, 0.412, 0.327, and 0.482 respectively.

Note that the NOS scores in the main body of the paper (using ANTs) are higher than these (using FSL), indicating better performance of ANTs at skull base registration in this dataset. However, these supplementary results are similar to the results in the main body of the paper, verifying the trends in the data and confirming that our results are not particularly sensitive to the choice of registration algorithm.

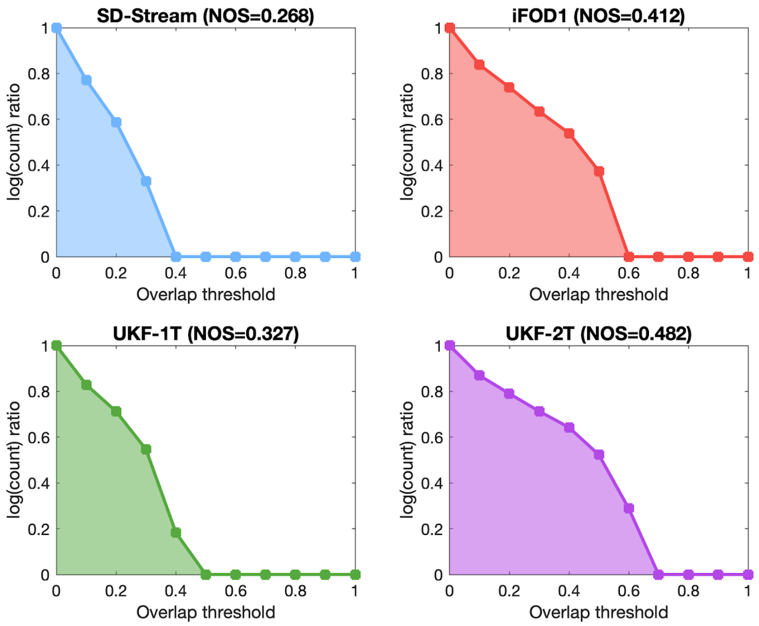


Figure S4 NOS of conjunction images generated by different tractography methods. The y-axis shows the log ratio (the ratio that is summed in equation 1), while the x-axis shows different threshold values of conjunction images.

**Supplementary material 6: wDice for each subdivision**

In addition to the inter-expert reliability validation on the entire RGVP reconstructed for each tractography method (Section 3.5 in the main paper), we also performed an experiment to assess the reliability for each RGVP subdivision. For each tractography algorithm and each subject under study, we computed the weighted Dice (wDice) score (Cousineau et al., 2017a) to measure the spatial overlap of each RGVP subdivision selected by the two different experts (G.X. and S.Y.). We note that for each tractography algorithm, we only included the RGVP subdivisions that were able to be identified by both experts. The results show that different tractography methods have different preferences, as shown in Table S2. The iFOD1 and UKF-2T methods have higher mean wDice scores in the decussating subdivisions than the non-decussating subdivisions, whereas UKF-1T has higher mean wDice scores in the non-decussating subdivisions than the decussating subdivisions. The iFOD1 and UKF-2T methods have a tendency to reconstruct decussating fibers, and the UKF-1T method tends to reconstruct non-decussating fibers. This conclusion is similar to the experiment of the percentage of decussating fibers.

| Supplementary Table 2: wDice scores for each RGVP subdivision and entire RGVP wDice score | | | | | |
|---|---|---|---|---|---|
| | LL | LR | RL | RR | Entire RGVP |
| SD-Stream | 0.76±0.25 | 0.80±0.18 | 0.77±0.29 | 0.79±0.22 | 0.74±0.19 |
| iFOD1 | 0.69±0.22 | 0.69±0.18 | 0.73±0.13 | 0.64±0.25 | 0.74±0.16 |
| UKF-1T | 0.86±0.15 | 0.77±0.24 | 0.78±0.15 | 0.81±0.23 | 0.82±0.18 |
| UKF-2T | 0.66±0.18 | 0.74±0.16 | 0.80±0.11 | 0.70±0.18 | 0.81±0.09 |
